# Optimized Biomechanical Design of a Pulsatile Fontan Conduit for Congenital Heart Palliation

**DOI:** 10.1101/2024.06.19.599796

**Authors:** Nir Emuna, Alison L. Marsden, Jay D. Humphrey

## Abstract

The evolution of palliative surgical procedures for children born with congenital heart defects has proven remarkably successful in extending life, but the resulting non-physiological circulation predisposes to myriad sequelae that compromise quality of life and overall life span. Among these procedures, standard-of-care Fontan completion surgery bypasses the nonfunctional ventricle and provides steady flow of deoxygenated blood to the lungs via a synthetic conduit that typically connects the inferior vena cava to a pulmonary artery. This altered circulation reduces cardiac output, elevates central venous pressures, and possibly contributes to adverse remodeling of the pulmonary vessels. There is, therefore, strong motivation to develop a next generation Fontan conduit capable of serving as a sub-pulmonic pulsatile pump, and there are now several reports of initial attempts. None of these studies have been driven by biomechanical considerations, however, and none have achieved the desired functionality. We thus present a novel analytical framework to improve design and guide fabrication by focusing on the microstructure and material properties of the contractile myofibers and associated passive matrix. Our optimized designs simultaneously ensure desired levels of stroke volume, ejection fraction, and pressure generation given constraints on Frank-Starling myofiber contraction and the limited space within the thoracic cavity of a three-to four-year-old child. This analysis also highlights the need to minimize any associated axial force or torque generation that a pulsatile conduit could transmit to the host vessels at the requisite anastomoses.

## INTRODUCTION

Congenital heart defects (CHDs) are the most common birth defects (approximately 8-10 out of every 1000 births), and among the most devastating. Remarkable advances in pediatric cardiac surgery over many decades – first ligation of a patent ductus arteriosus in 1938, first Blalock-Taussig shunt in 1944, first report of the Glenn procedure in 1958, first report of the Fontan procedure in 1971, and first report of the Norwood procedure in 1981, each with subsequent modifications – have enabled most children born with CHDs to live into adulthood. Yet, these individuals are at higher risk for early onset sequelae that affect multiple organs, including the heart, intestine, liver, and lungs (de Leval and Deanfield, 2010; Allen et al., 2017; Alsaied et al., 2020). There is, therefore, a pressing need for increased understanding and improved treatments.

Amongst these palliative surgical procedures, Fontan completion is typically performed when the affected child is two-to-four years of age. The extra-cardiac approach involves connecting the inferior vena cava (IVC) directly to the pulmonary arteries to bypass the nonfunctioning ventricle and deliver deoxygenated blood to the lungs. The current standard-of-care Fontan conduit is a cylindrical PTFE (GoreTex) graft. Albeit life-extending, this Fontan procedure results in a circulatory physiology characterized by reduced cardiac output and significantly increased central venous pressure, both of which likely contribute to early onset sequelae (Rychik et al., 2019; Kutty et al., 2020). For this reason, several groups are investigating alternate strategies that include sub-pulmonic mechanical pumps. Of importance herein, other groups are exploring regenerative medicine approaches to develop a pulsatile conduit to replace the passive GoreTex conduit with the hope of lowering central venous pressure and reducing sequelae. To date, however, none of the reported tissue engineered pulsatile conduits generate sufficient pulsatile pressures; maximum reported values range from 0.1 to 0.8 mmHg (Seta et al., 2017; Tsuruyama et al., 2019; Park et al., 2020; Endo et al., 2022; Köhne et al., 2022), well less than the ∼25 mmHg generated by the right ventricle. Clearly, the success of a pulsatile Fontan conduit will depend primarily on the availability of large numbers of robust cardiomyocytes needed to achieve the desired pulsatile blood pressure and flow. Yet, other factors will play critical roles, including optimized conduit dimensions, an extracellular matrix stiffness and anisotropy that facilitates diastolic filling, cardiac myofiber orientation and electrophysiological coordination, vascularization, and mechanobiologically favorable interactions with the host tissue at the sites of anastomosis.

We present a novel analytically based biomechanical analysis of a pulsatile conduit for Fontan completion as a step toward identifying an optimal design. An ideal surgical correction would effectively replace the critical function of the right heart – that is, yield appropriate stroke volumes while converting a steady, low-pressure venous return at ∼3 mmHg into a pulsatile output to the pulmonary arteries at 20-30 mmHg. Moreover, such a conduit must not overly stress the host IVC and pulmonary artery at the sites of anastomosis (Ban and Humphrey, 2023). High mechanical stresses can induce adverse mechanobiological responses or catastrophic mechanical failure. Thus, although not previously considered, multiple mechanical factors must be optimized simultaneously to achieve the desired performance. Of course, a pulsatile conduit will need to accommodate cardiovascular development throughout childhood, namely, changing dimensions, properties, and demands on cardiovascular and cardiopulmonary tissues due to somatic growth (Sluysmans and Cohen, 1985; Ochiai et al., 2010; Kathuria et al., 2015). We restrict our attention herein to the initial design, however, with geometrical considerations influenced strongly by the size and shape of current standard-of-care conduits.

## MOTIVATION and METHODS

### Geometric Constraints

Although subsequent derivations are general, or can be generalized, for purposes of illustration we used representative values and ranges of key cardiovascular parameters to define an initial design space for Fontan completion in children three to four years old. These parameters include the desired stroke volume (SV), ejection fraction (EF), and Frank-Starling cardiac myofiber shortening (from stretch λ_*f*_) as well as anatomical restrictions on luminal diameter at end diastole (*d*) and, in the case of a cylindrical tube, diastolic length (*l*). For example, a four-year-old child has a heart rate (HR) ∼103 bpm, right ventricular (RV) systolic/diastolic pressure of ∼27/7 mmHg, RV end diastolic volume (EDV) of ∼50 ml, and EF of ∼0.64 (Graham et al., 1973), where SV = EDV – ESV (end systolic volume), ejection fraction EF = SV / EDV, and cardiac output CO = SV_*_HR. For reference, the diameter of current standard-of-care Fontan conduits is 16-24 mm (Lee et al., 2007) and length is 40-60 mm (Alexi-Meskishvili et al., 2000; Ochiai et al., 2010). See Supplemental **Table S1** for representative ranges for multiple parameters relevant to early childhood (Lu et al., 2008; Cattermole et al., 2010; Kathuria et al., 2015).

Candidate geometries for a pulsatile conduit range from analytically complex (e.g., ellipsoidal, partial toroidal, or prolate spheroidal) to simple (spherical or cylindrical). It proves convenient first to consider the limiting cases of a sphere and cylinder, with the latter describing current Fontan conduits. Let the end diastolic luminal diameter be *d* for both, whereby EDV = *πd*^3^/6 for a sphere and EDV = *πd*^2^*l*/4 for a cylinder, with *l* its end diastolic length. Because EDV=SV/EF, *d* = (6SV/*π*EF)^1/3^ for the sphere and *d* = (4SV/*πl*EF)^1/2^ for the cylindrical tube. Given the small volume available to the surgeon for placement of a pulsatile conduit, these two simple relations place strong restrictions on geometry given target values of SV and EF.

Consider illustrative examples for a SV = 17 ml (assuming a lower bound resting IVC inflow alone, which accounts for ∼53% of the total stroke volume of ∼32 ml for a three-to-four-year-old child, with SV ∼22.2(age)^0.3^; Salim et al., 1995; de Simone et al., 1997) while targeting a range of potentially allowable diastolic diameters from 15-25 mm and, in the case of a cylindrical tube, a range of diastolic lengths from 40-60 mm (Alexi-Meskishvili et al., 2000; Itatani et al., 2009), both for an EF ranging from 0.5 to an optimistic 0.8 (Hurwitz et al., 1984). **Figure 1** shows that such a geometry-restricted spherical pump (panel A) cannot achieve the desired range of EF. By contrast, a geometry-restricted cylindrical conduit (panel B) can achieve SV = 17 ml with an EF within the upper end of this range. Hence, we focus below on cylindrical conduits consistent with the current standard-of-care geometry. Given that prototype conduits will need to be tested in animal models, often mice and sheep (Drews et al., 2020), similar analyses are performed easily for these species. Representative values for total stroke volume, EF, IVC fractional flow, and end diastolic diameter and length are listed in **Table S2** for these species, and results for a cylindrical conduit are contrasted across species in Supplemental **Figure S1**. Importantly, all reference below to the quantity SV will similarly be for a pulsatile conduit with IVC inflow alone. Results for confluence designs that include superior vena cava and IVC inflows can be computed similarly.

**1.**
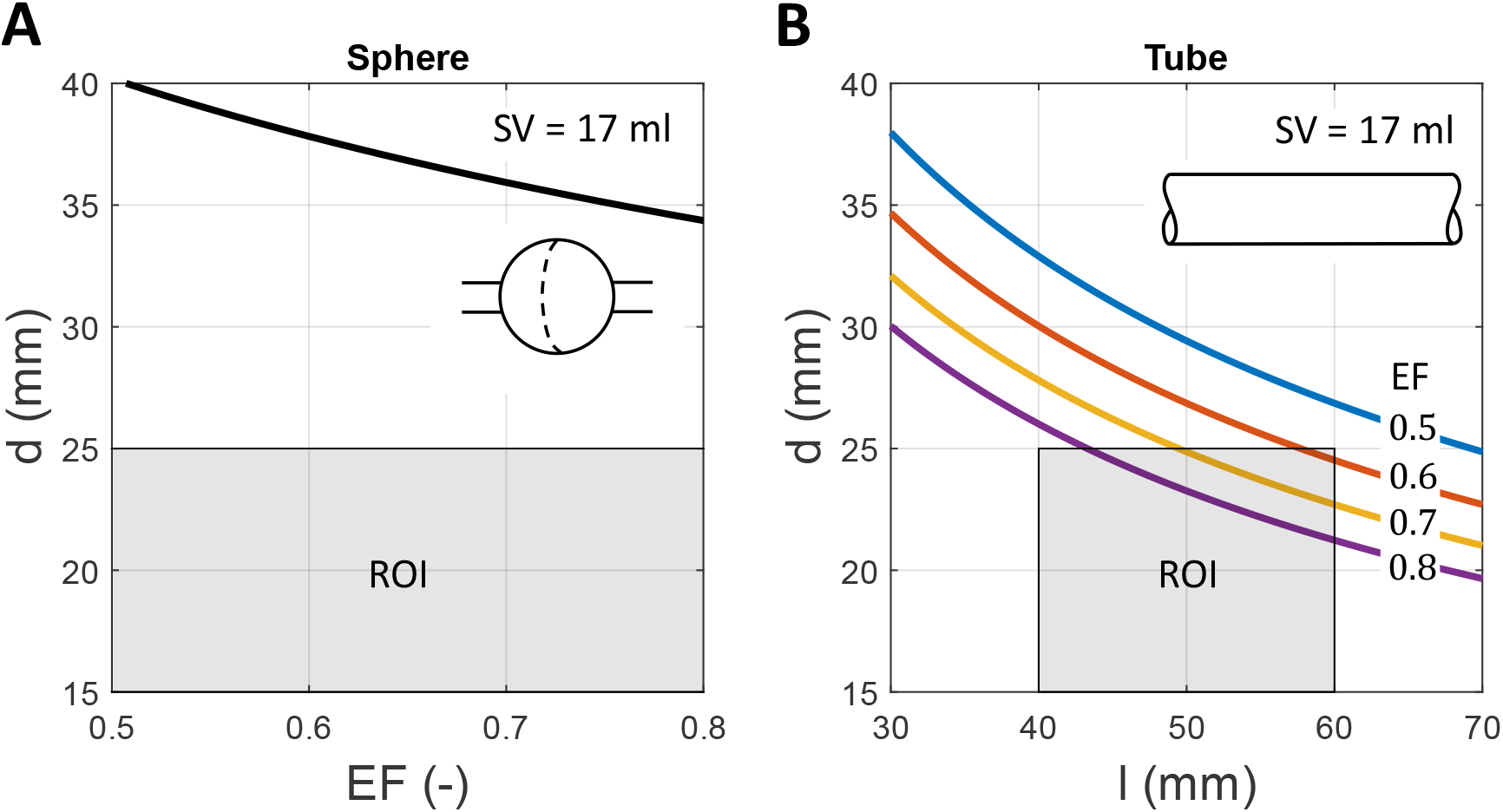
Basic conduit parameters for spherical (left) and cylindrical (right) geometries are deemed admissible if within the grey regions of interest (ROI) for an illustrative value of stroke volume (SV) equal to 17 ml and a range of ejection fraction (EF) from 0.5 to 0.8. The maximal available space in the thoracic cavity is assumed to be up to ∼25 mm in diameter based on the Abbott HeartMate 3 Left Ventricular Assist Device and up to ∼60 mm in length (if tubular), based on the distance between the intrapericardial part of the inferior vena cava and the undersurface of the right pulmonary artery for ∼3-year-old children (Alexi-Meskishvili et al., 2000). These acceptable dimensions are also consistent with current GoreTex Fontan conduits.

Another strong restriction is that maximal myofiber shortening is expected to range from 20-30% in the case of robust contractility. The EF for a cylindrical tube can be written

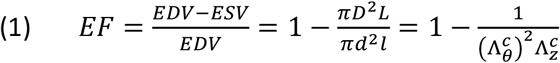

where stretch ratios 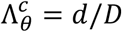 (ratio of luminal diameters) and 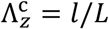 (ratio of axial lengths) highlight possible circumferential and axial stretches at the luminal surface due to passive filling of a valved conduit during each pumping cycle, from which isovolumic contractions can arise via myofiber shortening. As expected, changes in circumference impact EF more than changes in length, but both contribute.

Possible myofiber orientations within a cylindrical tube include axial, circumferential, and a combination of the two. Considering an orthogonal combination, even robust biaxial contractions along circumferential and axial directions only yield an EF of either 0.42 (for Λ_*θ*_ = Λ_*z*_ = λ_*f*_ = 1.2; **Figure S2A**) or 0.54 (for Λ_*θ*_ = Λ_*z*_ = λ_*f*_ = 1.3; **Figure 2A**), where we assumed maximal myofiber shortening of 20 and 30%, respectively, from stretch λ_*f*_. This simple result suggested the need to consider possible diagonal orientations of the myofibers, both symmetric and asymmetric, to achieve a higher EF given the geometric constraints noted above while reducing the need for excessive cyclic axial lengthening and shortening 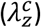 of the conduit over a pumping cycle. Of course, myofiber orientations exhibit a radially varying diagonal splay through the wall of a healthy heart that contributes to the twisting motion that augments ejection (Humphrey and Yin, 1989).

**2.**
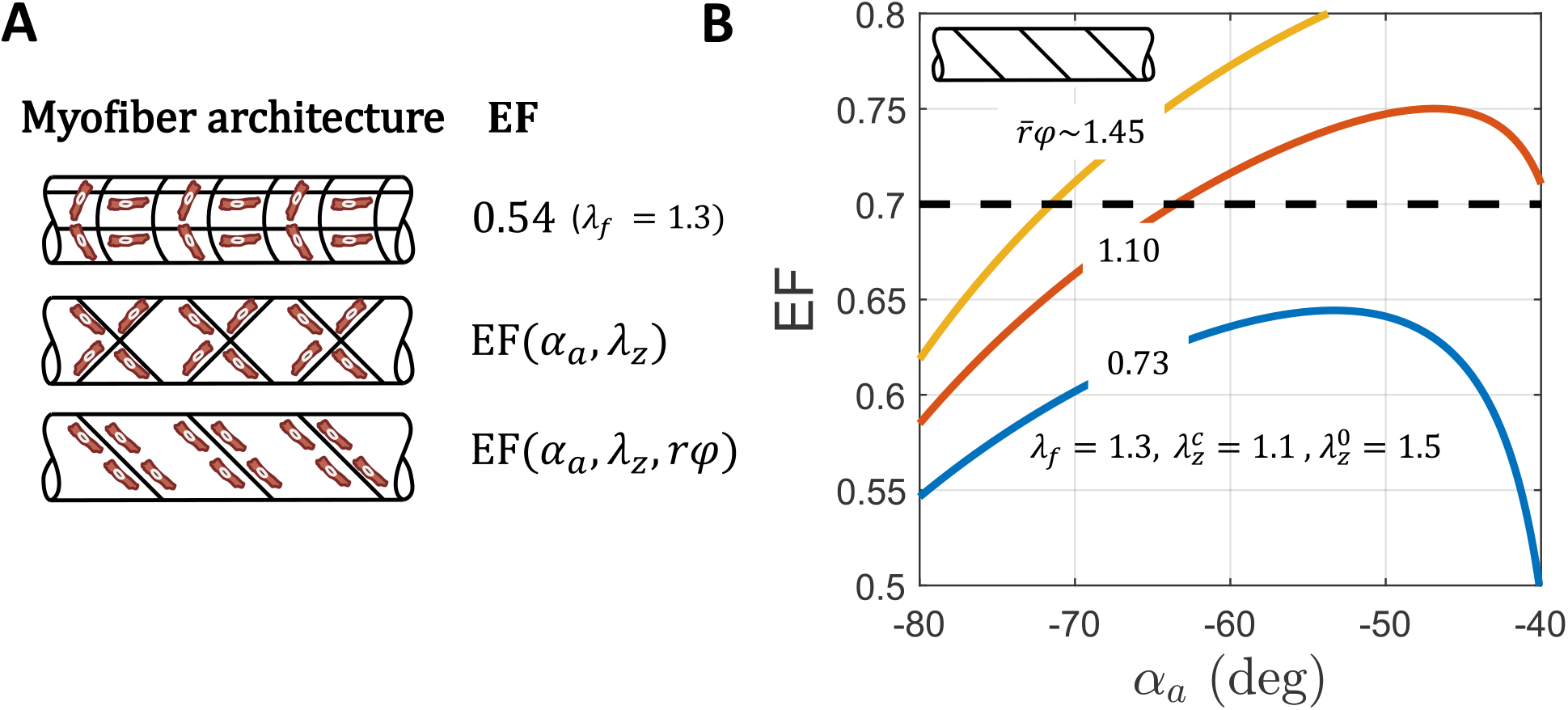
Blood is ejected via contraction of myofibers. Given a maximum active shortening of these filaments to be 20-30%, the ejection fraction (EF) is determined in large part by the orientation of the active filaments and allowable changes in overall dimensions (Sallin, 1969). Shown are calculations of EF as a function of fiber orientation (*α*_*a*_) for illustrative values of fiber shortening (λ_*f*_=1.3) and axial stretching during diastolic filling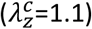, including the influence of twist in the case of an asymmetric orientation. See Figure S2 for similar calculations for λ_*f*_=1.2 and 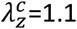.

### Finite Deformations

Consider a finite deformation of a cylindrical conduit that includes possible distension, extension, and torsion. The requisite passive matrix will control the pressure-induced distension and extension during passive filling and, in the case of asymmetric fiber orientations, an induced twist (cf. Emuna and Cohen, 2020). Active myofibers will then need to generate pressure during isovolumetric contraction (in a presumed bi-valved conduit; cf. Sharifi et al., 2021) and ejection, with a minimally pressurized reference configuration taken for convenience at the end of isovolumic relaxation (**Figure 3A**).

**3.**
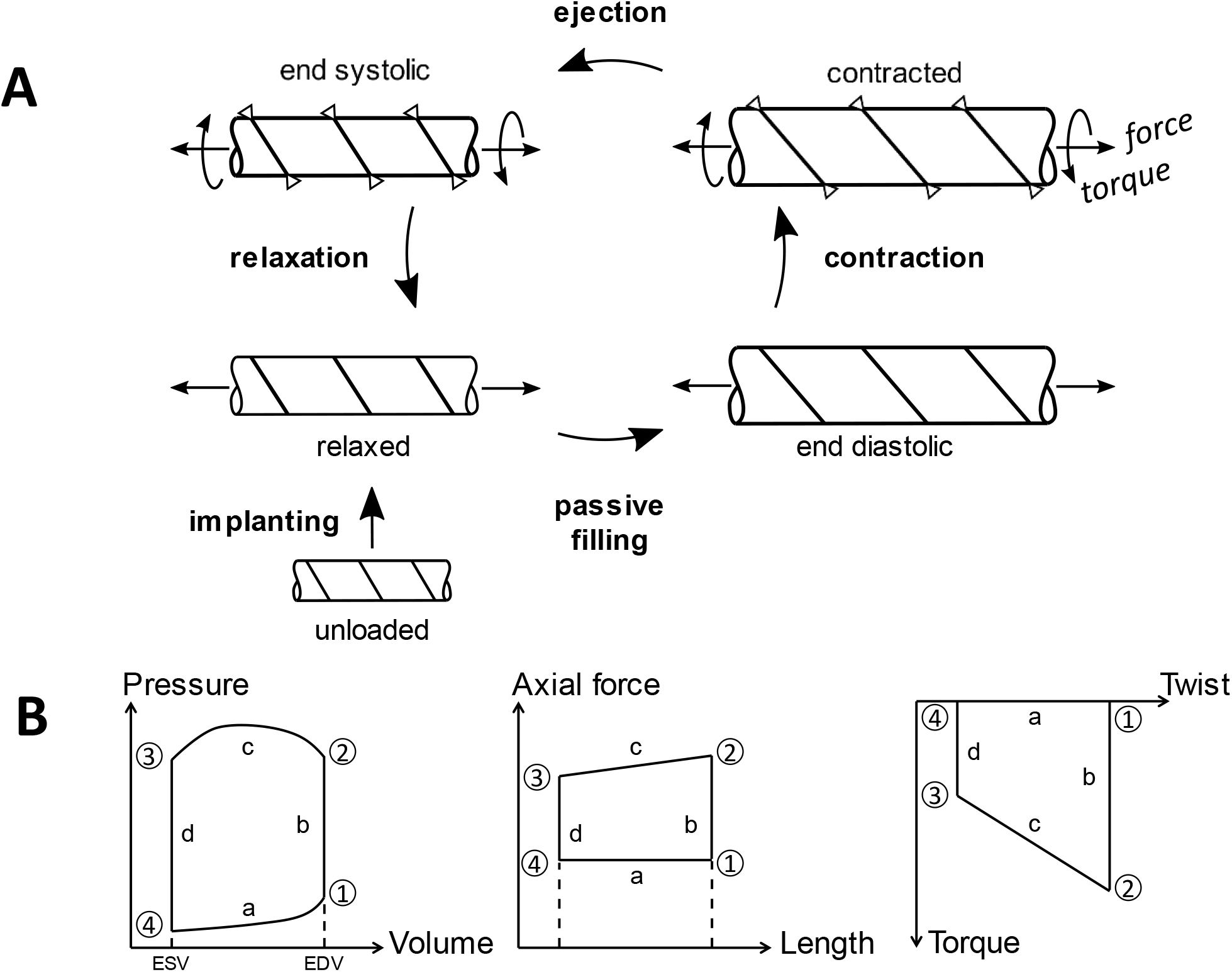
(A) Schema of a canonical pressure-volume loop biomimicking the right ventricle as well as associated states that a cylindrical pulsatile conduit will need to assume to replace ventricular function. (B) Illustrative pressure-volume, axial force-length, and torque-twist loops for a pulsatile conduit. The overall shape of a pressure-volume loop is defined by prescribed values of pressures (P1, P2, P3, and P4) and volumes (EDV, ESV) that are defined by physiological requirements (SV and EF). In contrast, shapes of force-length and torque-twist loops can be controlled by design parameters to optimize the overall function of the conduit.

Assuming that residual stresses are not introduced during fabrication of the conduit (i.e., absence of residual-stress related opening angles; Humphrey, 2002), let a material particle located at (*R, Θ, Z*) in a reference configuration be mapped to (*r, θ, z*) in a current (loaded) configuration via

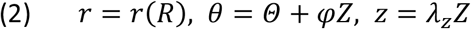

where *φ* = Φ/*L*_0_ is an angle of twist per unit unloaded reference length *L*_0_ and λ_*z*_ = *l*/*L*_0_ is an axial stretch ratio, noting that a cylindrical conduit may be pre-stretched during implantation (**Figure 3A**). Assuming that motions remain isochoric (i.e., volume of the conduit is preserved during deformation), the associated deformation gradient can be written in terms of physical components as (Humphrey, 2002)

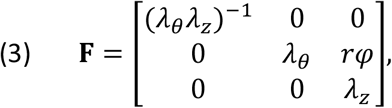

where λ_*θ*_ = *r*/*R*_0_ for any radial location within the wall. The associated right and left Cauchy–Green (deformation) tensors are, respectively,

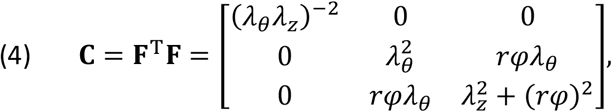

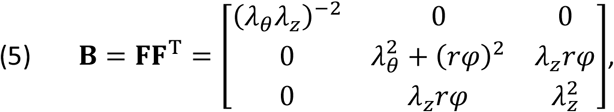

both of which are symmetric and, via the polar decomposition theorem, properly insensitive to rigid body motions (which is important given possible twist). It will prove useful below in constitutive modeling to note that the first principal invariant of **C** is

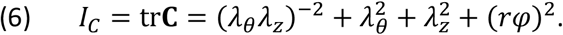

Next, let a unit vector along the direction of either a diagonal myofiber (defined by angle *α*_*a*_) or a passive matrix fiber (*α*_*p*_) within a circumferential-axial plane in the reference configuration be

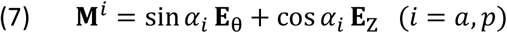

with (**E**_R_,**E**_θ_,**E**_Z_) a local referential orthonormal bases and angle *α*_*i*_ defined relative to the axial direction of the cylindrical conduit for any of the possible fiber families *i*, including those defining a transmural splay as in the normal heart (Humphrey and Yin, 1989). This unit vector can be mapped to a deformed configuration by

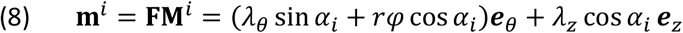

where (**e**_r_,**e**_θ_,**e**_z_) are local spatial orthonormal bases. The stretch 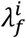 of any fiber initially oriented in direction **M**^*i*^ is thus given in terms of a coordinate invariant measure

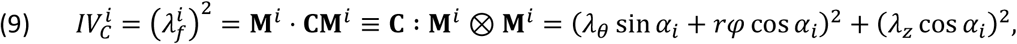

again with *i* = *a* for a myofiber capable of active contraction and *i* = *p* for a passive fiber, often collagen. For the former, this myofiber stretch defines the value from which it can shorten actively and generate force. In the absence of twist (i.e., *rφ* = 0), this relation recovers the special cases of myofiber extension equaling either axial (when *α*_*a*_ = 0) or circumferential (when *α*_*a*_ = *π*/2) stretch. This relation reveals further that twist-associated shear can alleviate some of the extension of an oriented myofiber as conduit-level stretches (λ_*θ*_, λ_*z*_) increase, as in diastolic filling. Another kinematic quantity that will prove useful below is ***m***^*i*^ ⊗ ***m***^*i*^ = **FM**^*i*^ ⊗ **M**^*i*^**F**^T^, namely

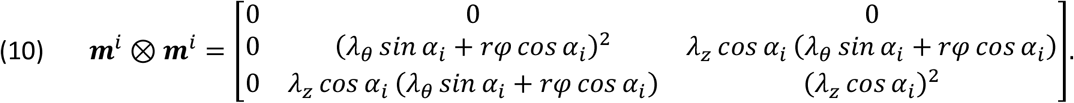

### Additional Motivating Results

Given the kinematics of distension, extension, and torsion, we can now contrast symmetric versus asymmetric myofiber architectures. Given a maximal myofiber shortening from stretch λ_*f*_ achieved during diastolic filling, the EF for both architectures can be expressed primarily in terms of myofiber angle (*α*_*a*_) and axial stretch (λ_*z*_), with the addition of shear (*rφ*) in the asymmetric case (**Figure S2A** and **Figure 2A**; see derivation in the Supplementary Materials). Note, here, that overall stretches can include contributions from possible pre-stretches induced at implantation and the cyclic stretches that arise over a pumping cycle, as, for example, 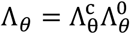 and 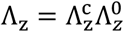 as overall circumferential and axial stretches at the luminal surface (i.e., at *r*/*R* = *a*/*A*), where superscripts (…)^0^ and (…)^*c*^ denote pre-stretch and cyclic stretch, respectively. Indeed, to preserve the natural in vivo axial stretch of the IVC, an ideal (compliance matching) conduit having a passive stiffness similar to that of the IVC would have a similar axial pre-stretch at implantation, which for illustrative purposes we let be 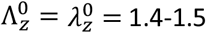 based on prior studies in sheep (Blum et al., 2022); for simplicity we assumed 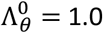 since luminal pressure will be essentially zero at the time of implantation. Assuming that the passive stiffness of most conduits will yet be higher than native, an axial pre-stretch of 1.5 serves as an upper bound.

For simplicity, first assume that the myofibers do not have an intrinsic deposition stretch at the beginning of filling. Preliminary results for symmetric myofiber orientations revealed strong limitations that prevent desired outcomes. For a maximal expected induced myofiber shortening from stretch λ_*f*_∼1.2-1.3, achieving a target EF would require the myofiber to be oriented towards the axial direction (**Figure S3A**), with high generated axial loads (**Figure S3B**) but low to unphysical (close to zero or negative) generated filling pressures (see Supplementary Materials). Moreover, the radially averaged (mean) circumferential stretch would need to be 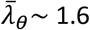 in this case, which is unlikely to be achievable.

Hence, focusing hereafter on asymmetric fiber orientations, **Figure S2B** shows examples for λ_*f*_ = 1.2 and 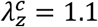 (due to diastolic filling) whereas **Figure 2B** shows examples for a more robust λ_*f*_ = 1.3 with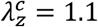, both with an upper bound implantation axial pre-stretch of 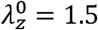 (and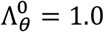).

Results are shown for different myofiber orientations running from nearly circumferential (*α*_*a*_ = -90 deg) toward axial (*α*_*a*_ → 0 deg) as well as for different values of shear. Given the goal of achieving values of EF equal to or greater than target values (e.g., dashed horizontal line in **Figure 2B**), it is seen that EF is dictated largely by the induced shear (or twist) that arises during passive filling given an asymmetric alignment of the passive fibers (e.g., collagen) within the matrix. Here, recall further that a tissue engineered pulsatile conduit having the current Fontan placement will be anastomosed to the IVC and a pulmonary artery, and by traction continuity any axial forces or torques (in the asymmetric case) generated by the contracting conduit will necessarily act on the host tissue (**Figure S4**). There is, therefore, a need for a detailed consideration of the constitutive relations and quasi-equilibrium equations.

### Constitutive Relations

Let the Cauchy stress **t** at any point within a conduit be determined by a sum of passive (matrix) and active (myofiber) contributions, namely, **t** = **t**^p^ + **t**^a^. Moreover, let these stresses be derived from scalar potential functions, with *W* = *W*^*p*^ + *W*^*a*^ (noting that the active part is not conservative, it is merely derived conveniently from a potential). Thus

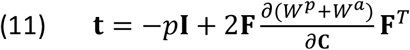

where the Lagrange multiplier *p* enforces isochoric motions, namely incompressibility. For illustrative purposes, let the passive stored energy be written as a combination of a neo-Hookean (isotropic, for a passive matrix) and a simple standard transversely isotropic (anisotropic, for a possibly asymmetric passive matrix fiber family, often collagen) relation (**Figure S5A**)

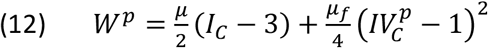

where *μ* (amorphous material) and *μ*_*f*_ (oriented material, or fibers) are passive material parameters. We further assume that the mechanical behavior of an active myofiber can be described by (Rachev and Hayashi, 1999)

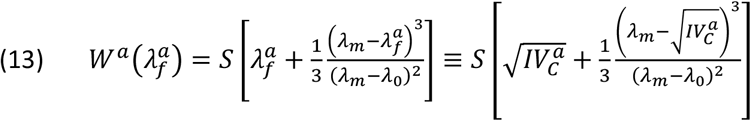

which yields a Frank-Starling type behavior (**Figure S5B**). The parameter *S* associates with the degree of muscle activation (with stress-like units) whereas 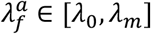 is the stretch of the myofiber with respect to the unloaded configuration of the conduit, with λ_*m*_ and λ_0_ values at which active force generation is maximal or ceases, respectively. One can also calculate the stretch of a myofiber relative to its individual unloaded length as λ_*F*_ = λ_*f*_/λ_0_ whereby 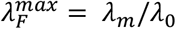 while 1/λ_0_ represents a type of deposition stretch of the active myofiber within the passive matrix. Values of λ_0_ < 1 allow the active myofiber to be pre-stretched within the nearly unloaded configuration of the conduit and thus to generate active stress in the end-systolic configuration.

Consequently, the total Cauchy stress (a symmetric tensor) is

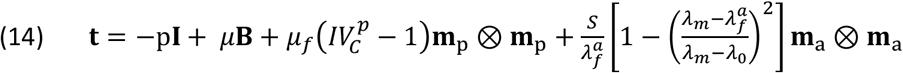

where, for convenience, the term in […] is denoted as the function 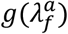 below. Note too, that we let 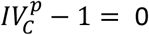 when 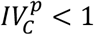 (e.g., passive bioprinted collagen fibers are assumed to not support compressive loads). Of course, the passive fibers and active myofibers need not be aligned with the same winding angle *α*. These equations are fully three-dimensional, and physical components of the Cauchy stress at any radial location within the wall are:

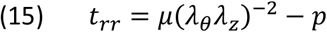

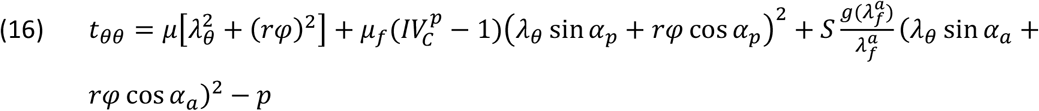

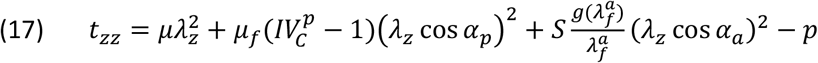

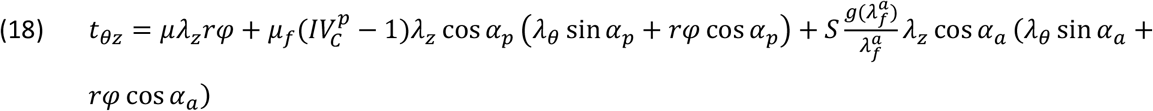

where in subscript/superscript *p* and *a* again denote passive fibers and active (contracting) myofibers, respectively. When the radial stress is small in comparison to the in-plane components, that is *t*_*rr*_ ≪ *t*_*θθ*_, *t*_*zz*_, this allows direct determination of the Lagrange multiplier *p*, namely

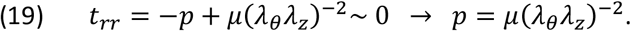

### Equilibrium

The three global equilibrium equations for finite distension, extension, and torsion of a cylindrical conduit require the following for the distending pressure *P*, axial force *f* (including a pressure term when proximal and distal valves close), and twisting moment (or torque) *T*

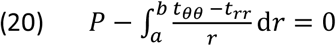

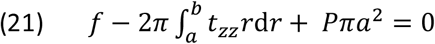

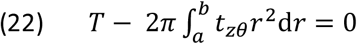

where *r* = *a* at the inner surface of the conduit and *r* = *b* at the outer surface, with *h* = *b* − *a* the deformed wall thickness. By contrast, on average, one need only require

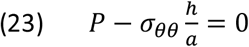

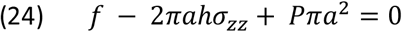

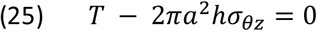

where *σ*_*θθ*_, *σ*_*zz*_, *σ*_*θz*_ are radially averaged (mean) values of *t*_*θθ*_, *t*_*zz*_, *t*_*θz*_ evaluated at the mid-wall value of circumferential stretch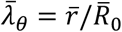, where 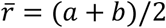 and 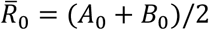 are average radii in current and reference configurations, respectively. The remaining deformation parameters are axial stretch λ_*z*_ = *l*/*L*_0_, where *l* and *L*_0_ are current and reference lengths, and twist per unit reference length *φ* = Φ/*L*_0_.

As noted above, conduit placement should include an implantation related axial pre-stretch in its relaxed state (with no myofiber contraction) to preserve the non-zero axial force in vivo in the host IVC (**Figure 3A**); perturbing the in vivo mechanical state of the IVC could lead to adverse post-operative remodeling. Consider, therefore, an axial pre-stretch to length 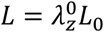 and possible pre-twist by an angle per unit reference length *φ*_4_ = Φ_4_/*L*_0_, with relaxed mid-wall radius 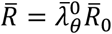 and thickness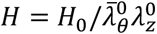. The superscript (…)^0^ again refers to the initial deformation during implantation whereas state 4 refers to the beginning of diastolic filling (see **Figure 3B**, which also shows state 1 at the end of diastolic filling, state 2 at the end of isovolumetric contraction, and state 3 at the end of ejection). Dimensions of the conduit at the end of diastolic filling (state 1, at EDV) are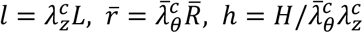, with *φ*_1_ = Φ_1_/*L*_0_ the twist angle per unit length. The superscript (…)^*c*^ again refers to cyclic deformations experienced by the conduit between ESV and EDV (due to filling) or between EDV and ESV (due to ejection). We can thus write the total circumferential and axial stretches between ESV (the reference state) and EDV (the maximum passively loaded state) as 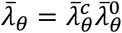 and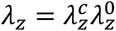, respectively. Mid-wall values of the square of the passive fiber stretch in the relaxed reference state (state 4, or ESV) are 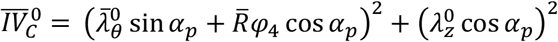 and similarly at the end of filling (state 1, or EDV) 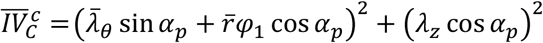. Mid-wall values of the square of the myofiber stretch at ESV are 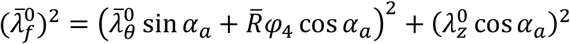 and at EDV are 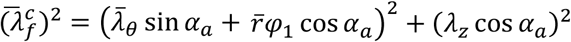.

Resultant loads in the relaxed state at the beginning of filling are pressure *P*_4_, axial pre-load *f*_4_ = *f*_0_, and zero torque (*T*_4_ = 0); for simplicity, we assumed that axial force and torque change little during passive filling given closure of only one valve, the low filling pressures, and lack of myofiber contraction. In the absence of data for the axial force in the IVC of a child, we used a measured physiological range of *f*_0_∼ 50-200 mN for sheep (Blum et al., 2022); this value is easily changed in the optimization algorithm when data become available.

### Structural Behaviors

Because a pulsatile conduit for Fontan completion should effectively replace the overall work done on the venous return by the right ventricle, its pressure-volume (P-V) behavior becomes important. **Figure 3B** illustrates four phases of P-V behavior for a cylindrical conduit that mimics that of the right ventricle: (a) passive filling during diastole (beginning with opening of the tricuspid valve), (b) isovolumic contraction (with the tricuspid valve and pulmonary valve closed), (c) ejection (beginning with the opening of the pulmonary valve), and (d) isovolumic relaxation. Whereas the heart is suspended and free to move within a lubricated pericardial sac, which minimizes reaction forces and moments, a current Fontan conduit is connected both distally and proximally via anastomoses with the IVC and pulmonary artery (**Figure S4**), which thus could support possible axial forces and torques. **Figure 3A** emphasizes that the work cycle of a pulsatile conduit is necessarily more complex than that in a standard-of-care GoreTex conduit. We posit that an optimized design should generate appropriate pressure– volume (P-V) behaviors while minimizing these two additional applied loads at the anastomoses, which requires consideration of associated axial force–length and torque–twist loops (which represent work done due to these additional loads).

To quantify the P-V loop in a cylindrical pulsatile conduit, it is convenient to satisfy equilibrium in four quasi-equilibrated states for the distending pressure (**Figure 3B**): *P*_1_ at the end of passive filling (with no myofiber contraction), *P*_2_ at the end of isovolumetric contraction with both valves closed, *P*_3_ at the end of ejection (with passive deformations coming only from placement pre-stretches) and *P*_4_ at the end of isovolumic relaxation (with effectively no contraction but possible deformation of the embedded myofiber due to the placement pre-load). For the radially averaged assumption, which facilitates intuitive understanding, (23) requires for the four successive states

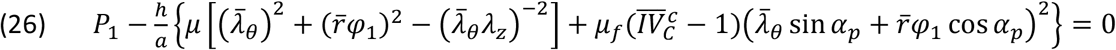

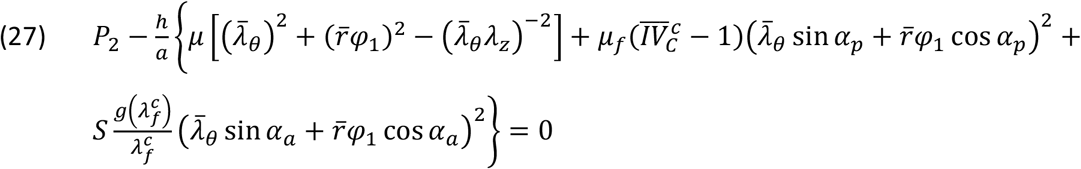

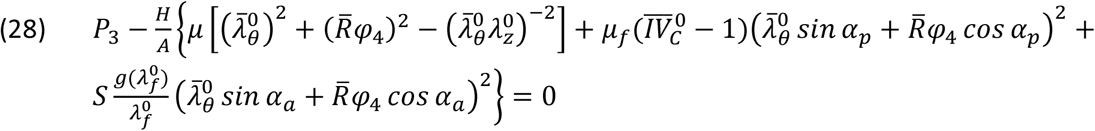

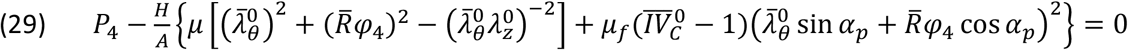

wherein we exploited kinematic conditions during isovolumic contraction and relaxation during which deformations are separately assumed to remain fixed (e.g., *φ*_1_∼*φ*_2_ and *φ*_3_∼*φ*_4_).

Whereas applied loads are often calculated in large deformation elasticity using a semi-inverse approach, that is, one finds the applied loads given the associated motions and material properties (Humphrey, 2002), we adopted an inverse-design approach. That is, we prescribed the desired primary loads of interest (e.g., pressures at different states *P*_1_, …, *P*_4_, axial pre-load *f*_0_) and associated deformations (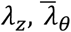, and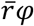, with the overbar again denoting a radially averaged value) that are consistent with the desired EF and SV (cf. **Figures 2B** and **S2B**) and then sought to identify those material properties (passive and active, including anisotropy arising from asymmetric fiber orientations) that enable this. Such an approach will guide the tissue engineer in selecting an appropriate matrix and myofiber, including orientation and contractile strength.

Toward this end, consider a relation between the shear induced during passive filling and the passive (collagen) fiber orientation. Generally, the associated twist could be obtained by determining λ_*z*_ and 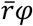 while satisfying overall axial force *f*∼ *f*_0_ and torque *T*∼0 (cf. Emuna and Cohen, 2021), with values of circumferential stretch varying from 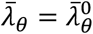 up to 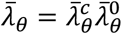, which is bounded by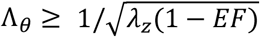. Here, however, we solve for the shear deformation 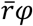 in two states as well as for the two material parameters *μ*_*f*_ and *μ*. The requisite equations are, for the passive filling phase (state 4 to state 1, ideally with *f*∼*f*_0_, *T*∼0 when *P* ∈ [*P*_4_, *P*_1_]), given by (24) and (25), namely, for axial force

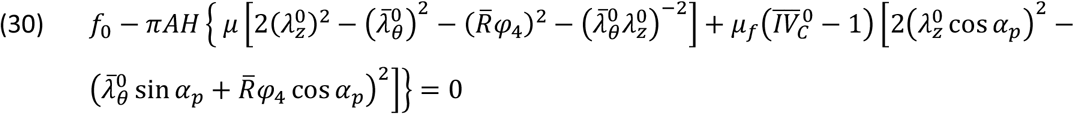

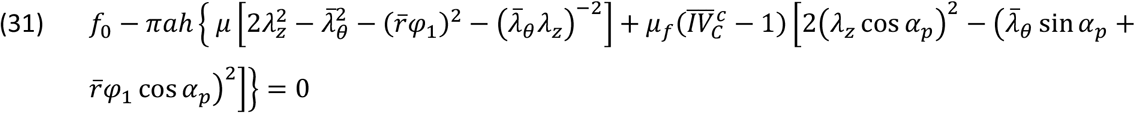

and for torque,

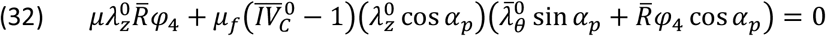

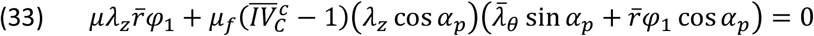

under the constraint 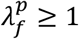 to ensure that the passive fibers are always under tension to prevent possible buckling while supporting the pressure-induced twist motion. The design problem is now fully defined and information necessary for design optimization is available.

### Parameter Sensitivity Studies

Solution of these 8 governing equations (26-29, 30-33) is sufficient to determine 8 unknowns 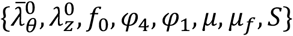 when the following parameters are prescribed or constrained {*P*_1_, *P*_2_, *P*_3_, *P*_4_, *a, h, A, H*} while others 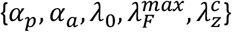[are varied parametrically for each solution. As discussed below, however, the entire range of possible fiber orientations from 0 to ±90 degrees for (*α*_*p*_, *α*_*a*_) need not yield admissible solutions. An admissible space can be mapped by solving the 8×8 system for given combinations of myofiber parameters 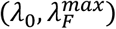 while scanning a range of preferred orientations, as, for example, [-50,-70] degrees. The results of these studies (discussed below) revealed that it is not straightforward to parametrically identify an optimal design and that there is a need to use formal methods of optimization.

### Design Optimization

Cardiac output (CO=SV_*_HR) is controlled differently in healthy children and those with a current standard-of-care Fontan circulation (Gewillig et al., 2010). For purposes herein, we prescribed SV as a hard constraint, that is, solutions can be repeated easily for any desired value of stroke volume as well as heart rate. We also constrained conduit radius and length to yield a volume at the end of diastolic filling that is less than or equal to that which is available to the surgeon in the chest cavity of a three-to-four-year-old child, which includes the volume occupied by current Fontan conduits. The primary focus, then, was to minimize the axial force and torque generated by the active myofibers during the pumping cycle while ensuring appropriate pressure generation to perfuse the pulmonary vasculature while respecting many different constraints (**Figure 4**). Toward this end, we simulated full axial force-length and torque-twist loops (recall **Figure 3**) and minimized the largest values of axial force and torque generated during myofiber contraction (isovolumic contraction or ejection). Although we did not assume that the maximum generated axial forces and torques necessarily emerged at the end of isovolumic contraction (state 2), it was nonetheless constructive to consider them. From (24) and (25), with the aid of (28) and (29), we find

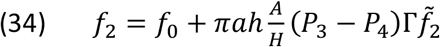

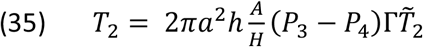

with normalized (unitless) axial force and torque defined as

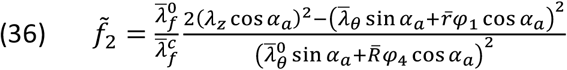

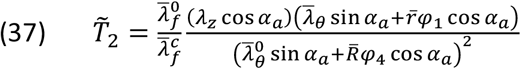

with 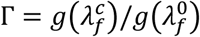 an effective, non-dimensional, measure of the contractility of the myofibers, combining the maximal stretch 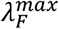 and deposition stretch λ_0_. Equations (34-37) reveal the relative influence of key parameters on the contraction-generated loads: geometry (*a, A, h, H*), physiology (*P*_3_, *P*_4_), myofiber properties (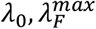 via Γ), deformation 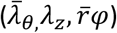 and microstructural orientations (*α*_*a*_, *α*_*p*_ via (36)-(37)).

**4.**
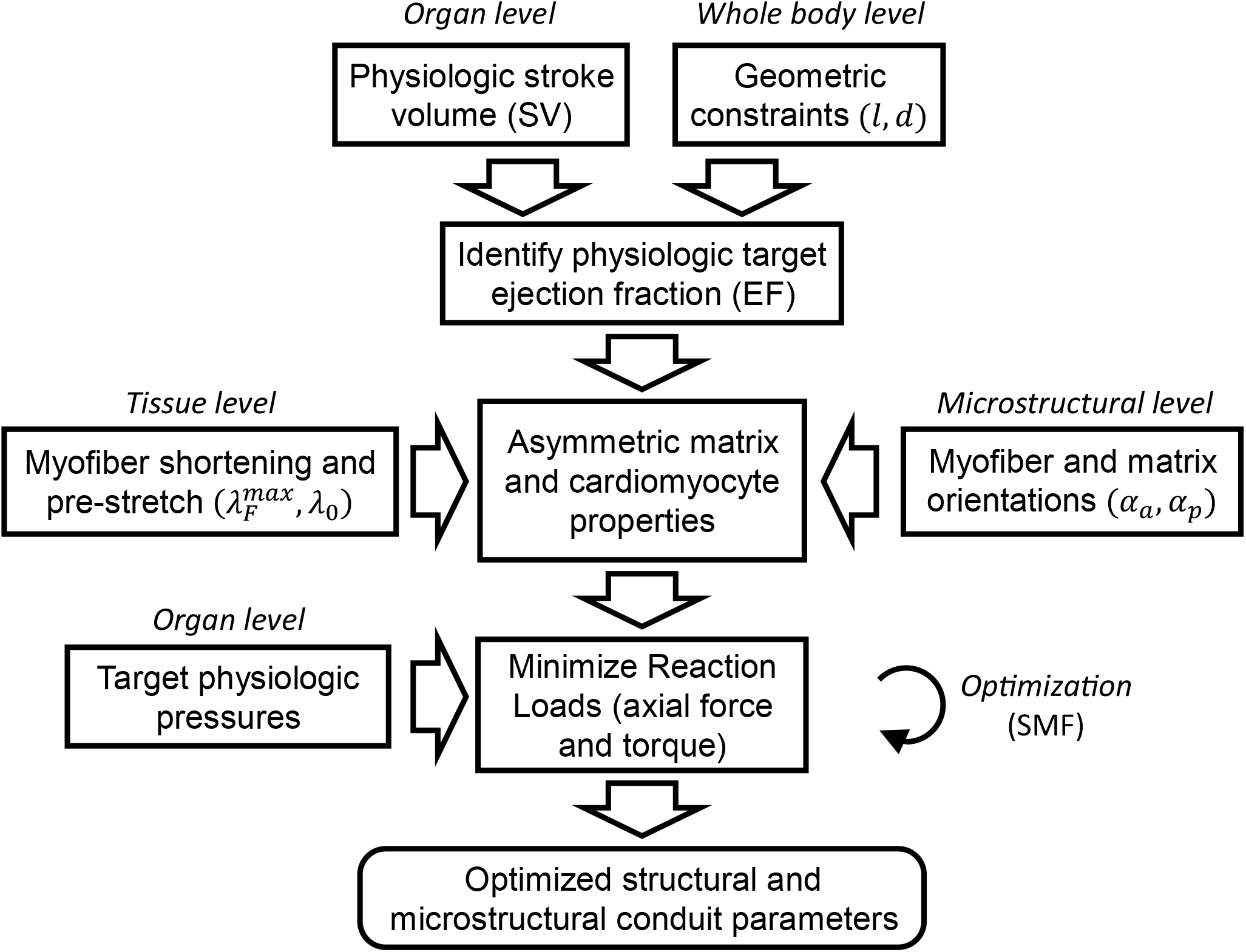
Suggested conceptual pipeline for optimization. Notwithstanding the simple cylindrical geometry considered, the many constraints and rich design space suggest that trial-and-error approaches are unlikely to be successful. The present pipeline enabled optimized solutions for a cylindrical pulsatile conduit based on minimization of the loads acting at the requisite anastomoses. Similar pipelines will be needed when expanding the search space to include 3D designs with transmural splays in fiber orientations, more complex non-cylindrical geometries, and future computational considerations that include fluid-solid-growth.

We now describe the optimization process step wise. For illustrative purposes, we prescribed as hard constraints the following: SV = 17 ml, EF = 0.7, P1 = 4 mmHg, P2 = 20 mmHg, P3 = 25 mmHg, P4 = 1 mmHg, l = 50 mm, H = 2 mm (see Supplementary Materials); although P1 and P4 will likely be higher in vivo, we used these values to consider an upper bound pressure generation during isovolumic contraction. Because the geometry and physiological function were either prescribed or bounded tightly, we sought first to optimize the deformation within an admissible range and so too contractile properties and alignment of the myofibers. Consider an objective function *J* that seeks to minimize the maximal loads *f* and *T* during the ejection phase. *J* can be weighted differently for the different contributors via *ρ* = *w*_*f*_/(*w*_*f*_ + *w*_*T*_). Hence, we sought to minimize

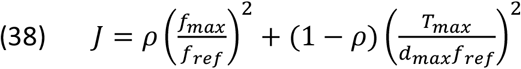

subject to multiple constraints: the passive fibers should remain under tension 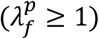 to support the induced twisting motion, the myofibers should remain under tension but not exceed the maximal stretch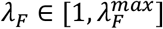, and the overall axial force should remain tensile (*f*_*min*_ > 0). Following many pilot simulations, we let weights for the two contributors be *w*_*f*_ = 1, *w*_*T*_ = 0.1, but other values could be considered. The need to weight the force more heavily than the torque was motivated by the functional form of the normalized loads 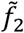 and 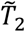 in (34)-(35). Specifically, 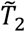 remains finite over a larger range of myofiber orientations, whereas 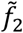 increases significantly beyond a threshold orientation (discussed below). Finally, note that *f*_*max*_ and *T*_*max*_ are assumed maximal absolute values of force and torque, *f*_*ref*_ is a reference value for force (assumed 100 mN), and *d*_*max*_ is the maximum loaded luminal diameter.

Free parameters to be optimized included: 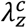 (the cyclic axial extension), λ_0_ (the deposition stretch of the myofiber at the unloaded configuration), 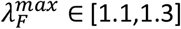 (the maximum stretch from which the myofiber can shorten fully), *α*_*a*_ ∈ [-50,-70] degrees (the asymmetric alignment of the myofibers relative to the axial direction), and *α*_*p*_ ∈ [-50,-70] degrees (the asymmetric alignment of passive fibers, if any), all subject to manufacturing limitations for material properties *μ* and *μ*_*f*_, with nondimensional parameter *γ*_*f*_ = *μ*_*f*_/*μ*. Restricting 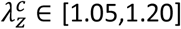 and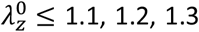, 1.2, 1.3 was based on the anticipated axial compliance of a bioprinted collagen matrix (∼10-30% elongation with respect to the unloaded state). Note that unloaded wall thickness *H* is determined independently from considerations of appropriate transmural values of wall stress (see Supplementary Materials).

The surrogate management framework (SMF) was selected for optimization. We used the Matlab code *surrogateopt* for constrained optimization. To ensure convergence to a global solution, we ran the code 100 times, each time with randomly generated initial guesses; each run was limited to 400 iterations. This approach was selected because SMF is a non-intrusive, derivative-free approach that has proven useful in diverse cardiovascular applications ranging from simulations of hemodynamics in the Fontan procedure (Yang et al., 2013) to parameterization of vascular growth and remodeling simulations (Sankaran et al., 2013) and the design of traditional (non-pulsatile) tissue engineered blood vessels (Szafron et al., 2019). The reader is referred to these papers for more details, noting that we also performed an independent gradient-based optimization that resulted in similar results (not shown).

## RESULTS

Figure 5. shows results from parameter sensitivity studies focusing on effects of changes in the orientations of the passive fibers and active myofibers (*α*_*p*_ and *α*_*a*_, respectively) given prescribed values of SV = 17 ml, EF = 0.7, H = 2 mm, and maximum end diastolic length of 50 mm (where the 25 mm diameter at end diastole derived from SV, EF, and l). In particular, Figure 5A shows effects of passive fiber orientation (via angle *α*_*p*_) on the mean shear 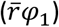 that is induced by the end of passive filling (i.e., state 1) as a function of three illustrative values of initial axial force (*f*_0_ = 50, 100, 200 mN) – the grey zone shows admissible values for which the passive fibers remain in tension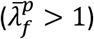) whereas the white zone excludes inadmissible values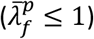. These results suggest, for relevant values of axial pre-load, that sufficient induced shear deformation can emerge for target values of SV and EF. This specific example was generated by solving equations (30)-(33) for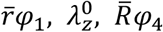, and *γ*_*f*_ for a range of passive fiber orientations *α*_*p*_ while prescribing representative values for *μ* = 1 kPa,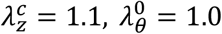. Additional results for this specific example are in **Figure S6**, with Panel A showing results for admissible values of the axial pre-stretch, again as a function of the initial axial force, which together impose a constraint on possible passive material properties (Panel B). A larger range is allowed for a passive fiber orientation up to *α*_*p*_ = -55, -60, -65 deg (for *f*_0_ = 50, 100, 200 mN, respectively).

Next, recalling the need to reduce both the axial force and torque generated during myofiber contraction, **Figure 5B** shows normalized loads (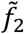 and 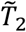 from equations (36) and (37), respectively) as a function of active myofiber orientation (via angle *α*_*a*_) for three admissible values of the passively induced shear 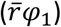 from **Figure 5A**, with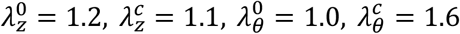. As it can be seen, axial force and the torque can each be minimized, but at different myofiber orientations. Notwithstanding intuition gained, this finding demonstrates the need to find a compromise design that would be difficult to isolate via parametric studies alone. That is, these results demonstrated the need for formal methods of optimization (equation 38) to achieve the best overall design. Note, too, that the actively generated force can become negative when the orientation of the myocytes is above certain threshold (towards circumferential), which is undesirable. Hence, the optimization must be constrained to yield physically desirable behaviors across multiple metrics.

**5.**
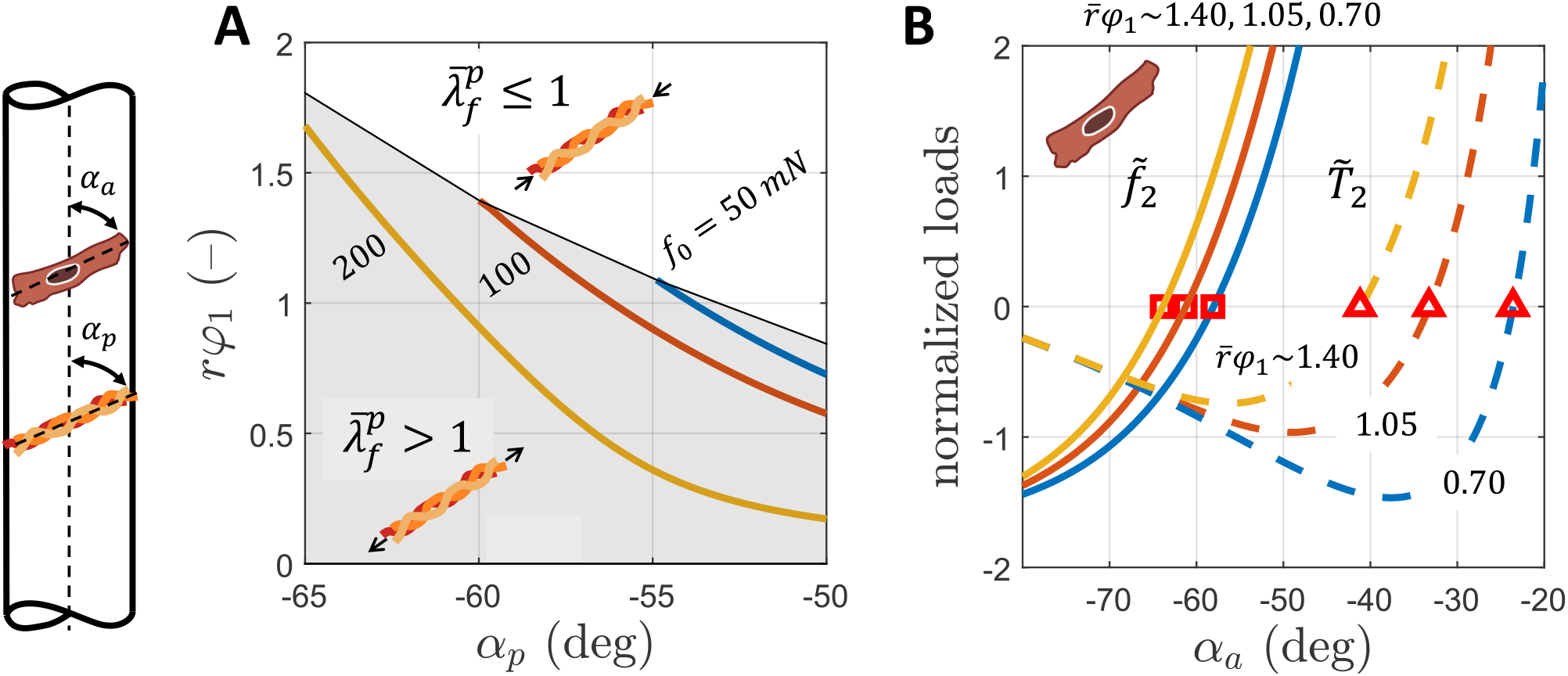
(A) The level of induced shear 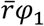 generated by asymmetric passive fibers at the end of diastolic filling as a function of fiber orientation *α*_*p*_ for three different values of admissible axial pre-load *f*_0_; admissible passive orientations *α*_*p*_ are found in the grey box. Values of parameters used to generate this panel include *μ* = 1 kPa,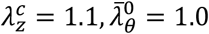; equations (30)-(33) that define the induced twist during passive filling were solved to generate the curves (without equations (26)-(29) that define the P-V loop). (B) Restricting attention to admissible values, generated normalized loads 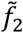 (force) and 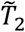 (torque) at the end of isovolumetric contraction as a function of active myofiber orientation *α*_*a*_ for increasing levels of shear 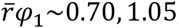, 1.05, and 1.40. The curves in this panel were generated by solving equations (36)-(37) with parameter values 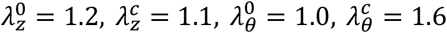, *r* = 12 mm, *L* = 30 mm, Φ = 100, 150, 200 d g, Φ_4_ = 20 d g.

Results from the Surrogate Management Framework are shown in **Figure S7** for one set of constrained optimized solutions for the special case of equal orientations of the passive fibers and active myofibers (*α*_*a*_ = *α*_*p*_), again with SV = 17 ml, EF = 0.7, H = 2 mm, and maximum end diastolic length of 50 mm (where again, the 25 mm diameter at the end diastole derives from SV, EF, and l). The resulting P-V loop (Panel A) respects prescribed values for P4 = 1 mmHg (end of isovolumic relaxation), P1 = 4 mmHg (end of passive filling), P2 = 20 mmHg (end of isovolumic contraction), and P3 = 25 mmHg (end of ejection), which are reasonable values for a pulsatile conduit with IVC inflow. The associated deformation during passive filling (states 4 to 1) reversed during ejection (states 2 to 3) with the assumed Frank-Starling type of behavior. Importantly, Panel B shows that isovolumetric contraction increased the axial pre-load from ∼120 mN up to ∼140 mN while this value decreased to ∼40 mN during mid-ejection and then back to ∼100 mN by the end of the ejection. Thus, the highest axial force was at the end of isovolumetric contraction and the lowest was during passive filling. All values remained within the desired physiological range based on values for the IVC in sheep (*f*_0_∼200 mN). Panel C shows, however, that a relatively high torque emerged at the end of isovolumetric contraction, ∼ -2.0×10^4 mm x mN. The associated range of twist corresponded to a change in angle along the length of the conduit from ∼0 to 100 degrees, hence the effect of pre-twist (Φ_4_) was small. Panel D provides details on changes in circumferential, myofiber, and axial stretches as a function of increasing pressure during passive filling. The range of myofiber stretch was ∼10-30% (i.e., the full range of myofiber stretch), with a myofiber deposition stretch of ∼10% corresponding to λ_0_ = 0.92. The cyclic circumferential stretch was within set values but required a significant dilatation during filling, up to 60% stretch with respect to the relaxed diameter (determined by EF and geometric constraints, see Eq. (1)); the optimized cyclic axial stretch was ∼20%. Both cyclic stretches are presented as normalized values with respect to the relaxed state. Finally, Panel E shows the induced angle of twist with increasing filling pressure, with a low pre-twist ∼10 deg at the beginning of passive filling and ∼100 deg at the end of passive filling.

Inasmuch as these results suggested that the desired pressure generation could be achieved with realistic constraints on geometry and myofiber orientation while generating modest axial forces and torques, there was further motivation to study the case of unequal passive fiber and active myofiber orientations (*α*_*p*_ ≠ *α*_*a*_). **Figure 6** shows optimized results similar to those in **Figure S7** except for unequal fiber orientations, which resulted in a broader parameter search space. Again, values of pressure were prescribed: P4 = 1 mmHg, P1 = 4 mmHg, P2 = 20 mmHg, and P3 = 25 mmHg for the same SV = 17 ml, EF = 0.7, and end diastolic geometry. Panel B shows the associated axial force-length loop with a pre-load of ∼60 mN. As it can be seen, axial load changed little during isovolumic contraction (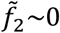 from equation (36)). In addition, axial load decreased to ∼0 mN during ejection but increased up to ∼80 mN and then back to ∼60 mN following relaxation to start the cycle again. Hence, as desired, the axial load remained within or below the expected physiological range. Panel C shows, relative to results in **Figure S7**, that the generated torque was smaller (maximum of ∼ -1.7×10^4 mm x mN), but the necessary range of twist angle was greater (∼20-140 deg) and the effect of pre-twist (Φ_4_) was larger. Panel D reveals reasonable ranges of deformations (circumferential, axial, and myofiber) during passive filling over the pressure range 1-4 mmHg. Specifically, the range of myofiber stretch was ∼10-24% (i.e., there was some reserve in myofiber stretch since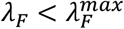), with a myofiber deposition stretch of ∼10% corresponding to 1/λ_0_ with λ_0_ = 0.92. The circumferential cyclic stretch was again within desired values of up to 60% with respect to the relaxed diameter (determined by the EF and geometric constraints, see Eq. 1) and the cyclic axial stretch was optimized at a lower value of ∼13%. Both cyclic stretches are presented as normalized values with respect to the relaxed state. Finally, Panel E shows the dependence of the induced twist angle on the pressure during filling, showing a moderate pre-twist ∼20 deg at beginning of passive filling and ∼140 deg at the end of passive filling.

**6.**
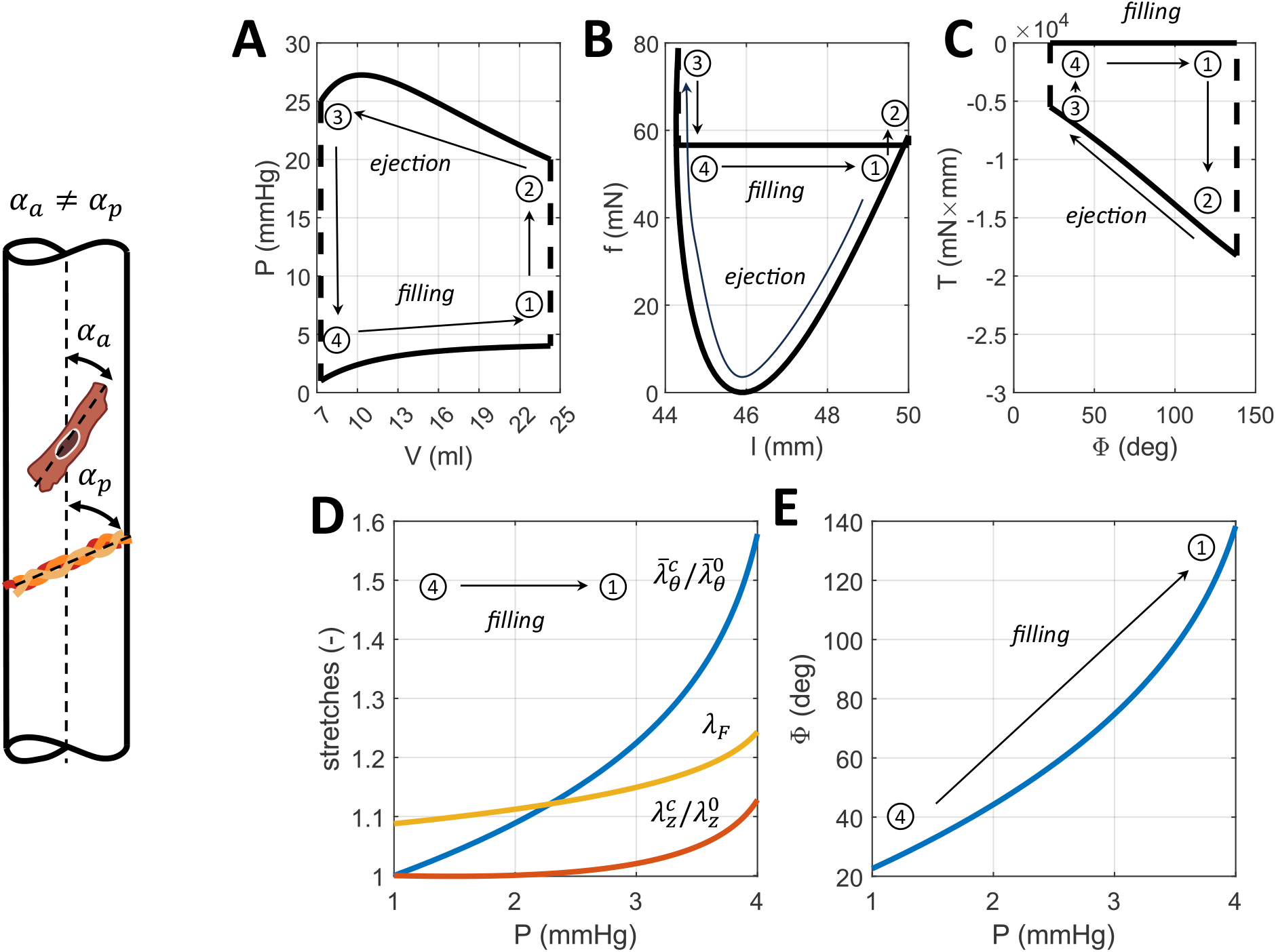
Illustrative (A) pressure-volume, (B) axial force-length, (C) torque-twist loops, (D) stretches during the passive filling phase, and (E) twist angle during passive filling for an optimized pulsatile conduit for the case of unequal active myofiber and passive collagen fiber orientations (*α*_*a*_ ≠ *α*_*p*_) for 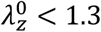 with *P*_2_= 20 mmHg, *P*_3_ = 25 mmHg, *P*_4_ = 1 mmHg, filling pressure *P*_1_ = 4 mmHg, EF = 0.7, and SV = 17 ml. See also Table 1.

**Table 1** lists values of many of the parameters that were optimized or prescribed in these simulations, both for equal (*α*_*a*_ = *α*_*p*_) and unequal (*α*_*a*_ ≠ *α*_*p*_) active and passive fiber orientations and for different values of allowable axial pre-stretches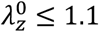, 1.2, 1.3. Note, for example, that the optimized results include values for matrix stiffness *μ* ∼1.5-1.7 kPa and active stress parameter *S*∼30 kPa, each independent of passive fiber orientation. **Table S3** lists illustrative optimized solutions across three species (human, sheep, mouse).

**1.** Optimized values of key design parameters for cases of either equal (*α*_*a*_ = *α*_*p*_) or unequal (*α*_*a*_ ≠ *α*_*p*_) orientations of active (a) myofibers and passive (p) fibers, each for different limiting values of axial pre-stretch (10 to 30%) that could be needed at the time of implantation to minimize adverse loading on the IVC. It is important to note that these specific values depend on the specific constraints, constitutive relations, and objective function used for illustrative purposes. Similar simulations testing different functions can now be easily performed as desired.

| | Parameter | $\alpha_a \neq \alpha_p$ | | | $\alpha_a = \alpha_p$ | | |
| --- | --- | --- | --- | --- | --- | --- | --- |
| | | $\lambda_z^0 < 1.1$ | $\lambda_z^0 < 1.2$ | $\lambda_z^0 < 1.3$ | $\lambda_z^0 < 1.1$ | $\lambda_z^0 < 1.2$ | $\lambda_z^0 < 1.3$ |
| Optimized parameters | $\alpha_a$ (deg) | -57 | -57 | **-59** | -56 | -60 | -63 |
| | $\alpha_p$ (deg) | -52 | -50 | **-53** | -56 | -60 | -63 |
| | $\lambda_F^{max}$ (—) | 1.27 | 1.30 | **1.26** | 1.30 | 1.30 | 1.30 |
| | $\lambda_0$ (—) | 0.92 | 0.91 | **0.92** | 0.93 | 0.92 | 0.92 |
| | $\lambda_z^c$ (—) | 1.14 | 1.12 | **1.13** | 1.20 | 1.20 | 1.20 |
| Preload and pre-stretch | $f_0$ (mN) | 44 | 47 | **57** | 47 | 89 | 121 |
| | $\lambda_z^0$ (—) | 1.10 | 1.10 | **1.13** | 1.10 | 1.20 | 1.29 |
| | $\bar{\lambda}_\theta^0$ (—) | 1.01 | 1.01 | **1.00** | 1.01 | 0.98 | 0.96 |
| Myofiber and passive stretch at end-diastole | $\lambda_F$ (—) | 1.25 | 1.28 | **1.24** | 1.30 | 1.30 | 1.30 |
| | $\lambda_f^p$ (—) | 1.01 | 1.01 | **1.01** | 1.01 | 1.01 | 1.01 |
| Material parameters | $\mu$ (kPa) | 1.56 | 1.60 | **1.51** | 1.70 | 1.59 | 1.49 |
| | $\gamma_f$ (—) | 7.4 | 7.3 | **8.9** | 2.6 | 3.3 | 3.9 |
| | $S$ (kPa) | 31.27 | 30.46 | **31.11** | 32.34 | 31.36 | 30.82 |
| Unloaded geometry | $L_0$ (mm) | 40.1 | 40.6 | **39.3** | 37.9 | 34.7 | 32.4 |
| | $R_0$ (mm) | 8.2 | 8.1 | **8.2** | 8.3 | 8.6 | 8.9 |
| | $H_0$ (mm) | 2.2 | 2.2 | **2.3** | 2.2 | 2.3 | 2.5 |
| Peak loads | $f_{max}$ (mN) | 71 | 93 | **79** | 242 | 175 | 144 |
| | $T_{max}$ (mN × mm) | 1.84x10<sup>4</sup> | 1.81x10<sup>4</sup> | **1.82x10<sup>4</sup>** | 1.99x10<sup>4</sup> | 1.90x10<sup>4</sup> | 1.84x10<sup>4</sup> |
| Objective function | $J$ (—) | 5.4583 | 5.6254 | **5.4554** | 11.1384 | 8.0649 | 6.8894 |

Although the optimized human solutions appeared to be physically reasonable for both equal (*α*_*a*_ = *α*_*p*_) and unequal (*α*_*a*_ ≠ *α*_*p*_) fiber orientations, it is prudent to examine the parameter space about the optimized values. **Figure S8** shows a mathematically admissible range of passive fiber and active myofiber orientations for different combinations of myofiber shortening 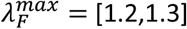 and deposition stretch λ_0_ = [0.90,0.95]. Clearly, more admissible solutions are possible with greater myofiber shortening and lower deposition stretch. Indeed, there were no solutions in the range of orientations [-70,-50] degrees for low shortening and high deposition stretch. Not all mathematically admissible solutions are physiologically relevant, however. **Figure 7** shows how particular constraints can limit the space of relevant solutions for diverse combinations of passive fiber and active myofiber orientations (the line of identity in each panel is for *α*_*a*_ = *α*_*p*_). Consider three constraints: (C1) *f*_*min*_ > 0, 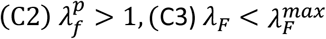. Consistent with **Figure S8**, more solutions are possible for high myofiber shortening and low deposition stretch (Panel B), less so for low myofiber shortening and low deposition stretch (Panel A), and the fewest for high myofiber shortening and deposition stretch (Panel C). Yet, the solutions in the more limited space in Panel C are the best optimized, with lower generated axial loads and torques and with larger range for myofiber shortening (**Table S4**). By contrast, the larger design space in Panel B would be more forgiving of fabrication errors in fiber orientations. Importantly, another advantage of the unequal orientations is clear – the space of relevant solutions along the line of identity (*α*_*a*_ = *α*_*p*_) is highly restricted. Indeed, the line of identity is outside of the space of relevant solutions in the case of the most optimized (lowest values of generated axial loads and torques) solutions in Panel C, meaning that only optimized solutions with different fiber angles can be considered for this combination of myofiber shortening and deposition stretch.

**7.**
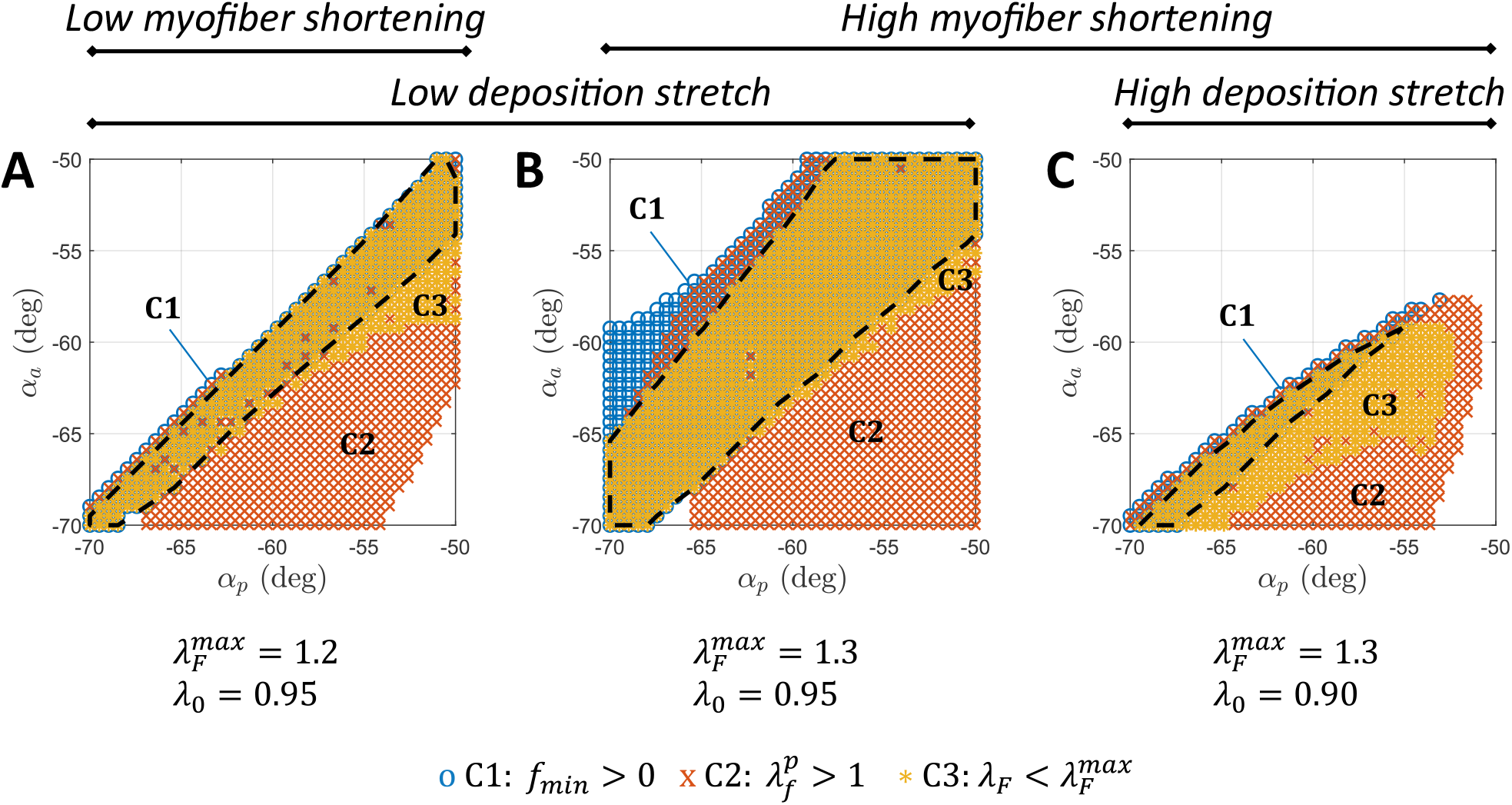
Design sensitivity maps. Each panel shows a mapping of admissible solutions over a range of passive fiber and active myofiber orientations (*α*_*p*_ and *α*_*a*_, respectively). Regions that respect three biomechanical constraints are highlighted: *f*_*min*_ > 0 (C1), 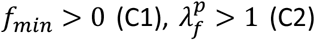, and 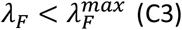. The union for the three constraints (dashed black curve) delineates the region of biomechanically admissible solutions. Values of parameters used to generate these maps are: 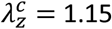, SV = 17 ml, l = 50 mm, H = 2 mm, EF = 0.7, *P* = 4 mmHg, *P*_2_ = 20 mmHg, *P*_3_ = 25 mmHg, *P*_4_ = 1 mmHg. See Table S4.

Up to this point, we have not addressed through-the-thickness variations of material properties and associated wall stresses in the conduit since we focused first on a radially averaged approximation to gain intuition in this highly parameterized and constrained nonlinear problem. This assumption yielded important analytical findings, including those highlighted in **Table 1** (i.e. average or mid-wall values of key parameters). Nevertheless, the range of validity and utility of this model are examined in **Figure S9**, with transmural distributions of wall stresses shown for key pressures along a P-V loop (recall Figure 3). Values of stress computed with the radially averaged and 3D models are shown in dashed and solid lines, respectively. As expected, radially averaged stresses generally capture mean values well. Yet, in the absence of fabricated residual stresses (cf. Szafron et al., 2017), the highest nonuniform wall stresses emerged at the highest isovolumic contraction pressure *P*_2_. Assuming that preferred levels of stress will be similar to those of the normal heart, ∼10-30 kPa (Burns et al., 1971), we anticipate a unloaded thickness of ∼2.4 to 2.9 mm, which if compared to a reference radius of ∼7.5-9.0 mm can be considered to be thick-walled in the absence of residual stresses (which otherwise tend to homogenize transmural distributions of stress; Humphrey, 2002). This finding suggests that a through-the-thickness splay of active myofibers would be beneficial, as in the native heart. An advantage of such splay is demonstrated qualitatively by comparing two distributions of active myofiber orientation (**Figure S10**): uniform versus a linear distribution with values *α*_*A*_ and *α*_*B*_ at the inner and outer surfaces, respectively. Shown, too, are associated distributions of myofiber stretch at the end of passive filling (before isovolumic contraction). The first has a linear distribution of stretch with values both above and below the threshold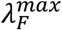. The second has a nearly uniform distribution of myofiber stretches, thus allowing all cardiomyocytes to contribute equally, thus suggesting that a linear splay can be beneficial and render the above results yet useful for design.

## DISCUSSION

Notwithstanding life-extending benefits of current standard-of-care GoreTex conduits for Fontan completion surgery, tremendous advances in regenerative medicine have led to an FDA-approved clinical trial of a next-generation tissue engineered vascular graft (TEVG) as a Fontan conduit (Drews et al., 2020; Breuer et al., 2023). Advantages of this and similar TEVGs are many, especially the more native structure and function, growth potential, and resolution of a chronic foreign body response, which may contribute to lower rates of calcification in Fontan TEVGs (Turner et al., 2024). Yet, these TEVGs have been designed as vascular, not cardiac, replacements; they are not designed to drive pulsatile flow to the pulmonary arteries at increased pressures, as achieved by the normal right heart. Loss of pulsatility via current Glenn and Fontan procedures may contribute to the reduced somatic growth of the pulmonary arteries post-Fontan completion (Ovroutski et al., 2009; de Leval and Deanfield, 2010; Shabanian et al., 2022) since cyclic loading may provide a unique mechanobiological stimulus for vascular growth, particularly during the early postnatal period (Rego et al., 2023). More importantly, absence of a sub-pulmonic pump in the current Fontan circulation results in a marked increase in central venous pressure, which likely contributes to the many sequelae, including Fontan associated liver disease and protein losing enteropathy (de Leval and Deanfield, 2010; Allen et al., 2017).

For these and related reasons, several groups have begun development of tissue engineered pulsatile conduits for Fontan completion (Biermann et al., 2016; Seta et al., 2017; Tsuruyama et al., 2019; Park et al., 2020; Endo et al., 2022; Köhne et al., 2022). Yet, none of these groups informed their design with theoretical or computational models to optimize design parameters. Rather, they appeared to use empirical trial-and-error approaches based on a tacit assumption that robust contractile cardiomyocytes will adapt to prescribed culture conditions and enable the necessary functionality. Interestingly, myofiber orientation has not been measured or emphasized in many of these studies though there have been observations of circumferential (Biermann et al., 2016), irregular (Seta et al., 2017), and axial (Köhne et al., 2022) orientations. Yet, as revealed by simple theoretical considerations herein, neither purely circumferential nor axial alignments, including combinations thereof, are likely to provide adequate function under known geometric constraints. Related to this, one study reporting a circumferential orientation implied a need “to orient [cardiomyocyte] fibers at specific angles” given the transmural splay of myofibers in the normal heart and citing a computational study of effects of fiber orientation on ventricular mechanics (Palit et al., 2015). Whereas computational models have long been used to design new candidate surgical approaches for Fontan completion (Marsden et al., 2010), to assess performance of the Fontan circulation post-operatively (Liang et al., 2014), and to predict possible surgical outcomes (Conover et al., 2018), all such studies have focused on the hemodynamics and thus computational fluid dynamics. There is, however, both regulatory motivation (Morrison et al., 2008) and ample evidence (Waters et al., 2021; Loerakker and Humphrey, 2022) that computational models can similarly be used to drive regenerative medicine solutions.

Focusing our attention on a possible cylindrical geometry, consistent with current GoreTex conduits and TEVGs, we developed a theoretical framework both to examine simple scenarios that build intuition and to optimize design parameters to generate adequate sub-pulmonic pulsatile pressures in the absence of a functioning right ventricle. We found, for example, that for the constraints considered, myofibers oriented symmetrically about the axis of the conduit would need to tend towards the axial direction to generate sufficient ejection fractions, but this would likely create excessive axial forces at the requisite anastomoses. Although asymmetric orientations towards the circumferential direction appear to be sufficient to generate appropriate perfusion pressures (∼20 mmHg) and stroke volumes (∼17 ml) for three-to four-year-old children, such orientations will generate both axial forces and torques at the anastomoses. Anastomosis geometry and function can be fundamental to overall success of a compliant conduit (Ban and Humphrey, 2023), hence it is surprising that this aspect of the design had not been considered heretofore. Parametric studies herein suggested that multiple orientations could yield modest values of actively generated axial force or torque while supplying sufficient perfusion pressures provided that the myofibers extended sufficiently during diastolic filling. That different values of myofiber orientation separately minimized the axial force and torque (Figure 5) revealed, however, that there was need formal optimization to minimize these loads simultaneously, thus yielding a weighted compromise solution. The associated optimized myofiber orientations and behaviors – achieved in part via passive twisting of the conduit during diastolic filling due to an anisotropic matrix – appears to improve function such that sufficient ejection fractions and stroke volumes may be realizable without requiring truly optimal myofiber shortening (of 30%), which is good news for those differentiating these cells. Given the design considerations and constraints imposed, recall that **Table 1** lists candidate optimized parameters for a three-to-four-year-old child for two basic cases: one preferred (*α*_*a*_ ≠ *α*_*p*_) and one that may be easier to fabricate (*α*_*a*_ = *α*_*p*_). Given that first-in-human studies will necessarily follow extensive animal studies (Drews et al., 2020), **Table S3** provides similar optimized parameters for both sheep and mice, which reveal similar design objectives across species.

Importantly, optimized solutions for a three-to-four-year-old child suggest slightly different orientations for the passive fibers (about -52 degrees) and active myofibers (about -58 degrees), thus favoring a slight circumferential orientation (that is, > 45 degrees from axial). They further suggest that there should be a modest axial pre-stretch in the conduit at implantation, ∼12%, and similarly an ∼12% axial stretch during each cycle of diastolic filling. That is, an asymmetric design will necessitate both distension and extension during passive filling to achieve target values of SV and EF. To facilitate ease of diastolic filling, the optimized solution suggested the need for a compliant underlying amorphous passive matrix, with stiffness of ∼1.5 kPa, though with oriented passive fibers that are about 9x stiffer than the matrix. These oriented passive fibers, most likely collagen, will enable the required twisting during filling that can augment reverse twisting during ejection when the myofibers contract. Critically, these active myofibers will need to generate a stress of ∼30 kPa and shorten about 25%, thus revealing the need for robust, coordinated, finely oriented cardiomyocytes. Finally, to achieve overall wall stresses comparable to those in the native heart, relaxed wall thickness will need to be on the order of 2-3 mm, which is much greater than the typical diffusion distance of oxygen in tissues, thus confirming the need for vascularization of the conduit itself. Since these vasa vasorum like vessels could also contribute to the stiffness and anisotropy of the conduit, their properties and orientation will need to be included within the functional optimization process. Murine conduits are expected to have the advantage of no need for vascularization, thus allowing early design iterations to focus first on optimizing the biomechanical behavior of the conduit itself.

Despite the insight gained, multiple limitations remain to be addressed. We did not consider non-cylindrical geometries, such as pear-shaped, which could prove to be superior to the simple cylindrical geometry studied. We did not address the need to ensure appropriate flow distributions to the left and right pulmonary arteries (Bachler et al., 2013; Yang et al., 2013), noting that this need not be part of the pulsatile conduit design but rather ensured via distal connectors to the pulmonary arteries; toward this end, we expect that passive only cuffs will be needed to suture the conduit to the host vessels, but we did not consider a functionally graded design. Indeed, we did not consider the possibility that the conduit may need to be segmented, with an atrium-like inlet to facilitate diastolic filling of a ventricle-like pressure generating conduit. We did not consider the electrophysiology and possible need for peristaltic versus pulsatile stimulation. We did not consider possible torsional-axial buckling (Hu et al., 2024) that may occur during the twist-driven ejection phase, noting that the torque levels during contraction may be above the stability thresholds of the conduit structure (cf. Emuna and Durban, 2020). Finally, we did not couple our optimization with either a wall growth and remodeling simulation (Szafron et al., 2019) to address the critical need to account for somatic growth or with a fluid-solid interaction simulation (Schwarz et al., 2021) to account for nonuniform distributions of flow- and pressure-induced mechano-stimuli to the cells within the conduit, which will necessarily need to include at the minimum endothelial cells to prevent thrombus, fibroblasts to maintain the matrix, and cardiomyocytes to generate the pressure. Of course, an integrated fluid-solid-growth model (Schwarz et al., 2023) would provide the most information. Indeed, there is significant concern that a pulsatile conduit having a Fontan-like placement may increase blood pressure in the superior vena cava (Hu et al., 2024), which could necessitate a different placement and possibly geometry. Many additional considerations are thus expected. Fundamentally, however, the present design considerations promise to guide all future design refinements, with the current analytical solution promising both to guide and to aid in verification and validation of future computational approaches that will be needed to address the many additional complexities, including overall impacts on the circulation within the lung, liver, and brain, often modeled using lumped parameter networks.

In summary, the present study shows that theoretical considerations can help to identify design goals and reduce the experimental search space (i.e., so that one can focus on theoretically viable designs alone), thus promising to accelerate research towards improved regenerative medicine solutions for CHDs. It was not surprising that our step wise approach from simple conceptual considerations to parametric studies to formal optimization led to a biomimetic solution – complementary passive and active anisotropy to increase SV and EF via a combination of extension, distension, and torsion similar to that of the native right ventricle. Indeed, just as in the ventricle, a transmural splay of myofibers appears necessary. Identifying specific target numerical values will be critical for final design, particularly since the optimal design space appears to be surprisingly narrow given tight constraints dictated by the geometry available to the surgeon and the requisite functional parameters, which for a Fontan conduit include adequate perfusion pressure, stroke volume, and ejection fraction while minimizing axial force and torque at the anastomoses.

## ACKNOWLEDGMENTS

This work was supported by a grant from Additional Ventures. Extensive conversations with and advice from Dr. Kirstie Keller, Vice-President of Additional Ventures, as well as other members of the Additional Ventures Cures Collaborative (Drs. Chris Breuer, Mike Davis, Nicole Dubois, Adam Feinberg, T-Y Hsai, Stacy Rentschler, Mark Skylar-Scott) are gratefully acknowledged. We also thank Dr. John Kelly for information on relevant cardiovascular parameters for sheep.

## SUPPLEMENTAL MATERIALS

### SUPPLEMENTAL METHODS

#### Expression of ejection fraction (EF) in terms of myofiber angle (α_a_) and the underlying deformation

The overall stretch includes contributions from both pre-stretch and cyclic stretch, with 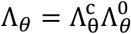 and 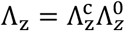 the overall circumferential and axial stretches evaluated at the luminal surface (*r*/*R* = *a*/*A*), where superscript (…)^*c*^ and (…)^0^ denote the cyclic (during a cardiac cycle) and pre-stretch (due to implantation) components, respectively.

##### 1) Symmetric diagonal fibers

The myofiber stretch evaluated at the luminal surface (λ_*θ*_ = *a*/*A* = Λ_*θ*_, λ_*z*_ = Λ_*z*_) is given by eq. (9) for a symmetric diagonal configuration in the absence of twist, namely

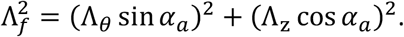

The expression of the overall luminal circumferential stretch is given by

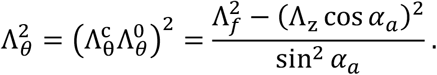

Hence, given the definition of ejection fraction (eq. 1)

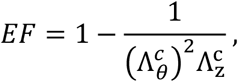

substitution of the expression for luminal cyclic circumferential stretch yields

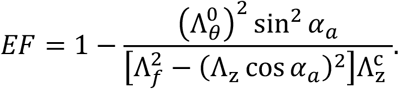

##### 2) Asymmetric diagonal fibers

The myofiber stretch evaluated at the luminal surface (*r* = *a*, λ_*θ*_ = *a*/*A* = Λ_*θ*_, λ_*z*_ = Λ_*z*_) is given by eq. (9) for an asymmetric diagonal configuration with twist, namely

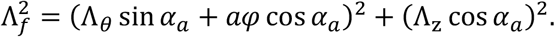

The expression of the overall luminal circumferential stretch is given by

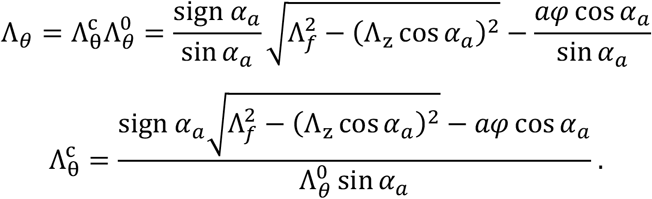

Hence, given the definition of the ejection fraction (eq. 1)

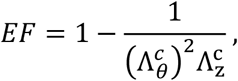

substitution of the expression for the luminal cyclic circumferential stretch yields

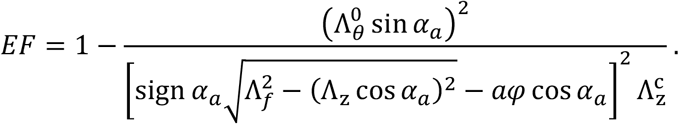

##### 3) Expression for the cyclic circumferential stretch for given EF evaluated at the luminal surface is simply

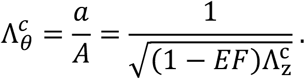

### Radially averaged symmetric conduit

For illustrative purposes, let the passive stored energy be written as a combination of a neo-Hookean (isotropic, for a passive matrix) and a simple standard transversely isotropic (anisotropic, for a possibly asymmetric passive fiber family) relation

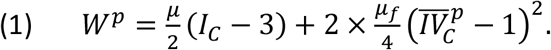

The mechanical behavior of the active myofibers is given by

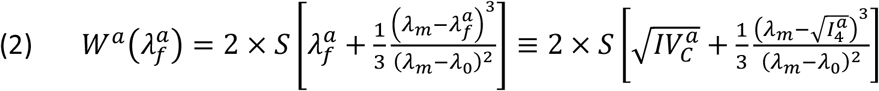

where the × 2 factor denotes the two symmetric diagonally orientated passive fibers (in *W*^*p*^) and myofibers (in *W*^*a*^).

The radially averaged stresses are

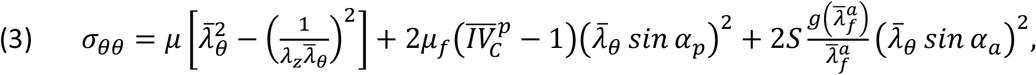

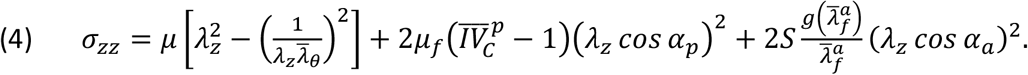

where an overbar denotes an evaluation at the mid-wall. Important kinematic quantities are

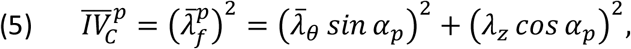

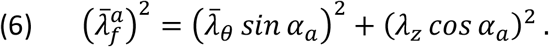

The distending pressure and axial force are given by global equilibrium,

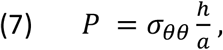

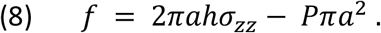

If we make the simplifying assumption that the axial force remains zero *f*∼0 (similarly, but identically with the asymmetric design where we considered an initial axial force *f*_0_ ≠ 0) during passive filling (*S* = 0), then

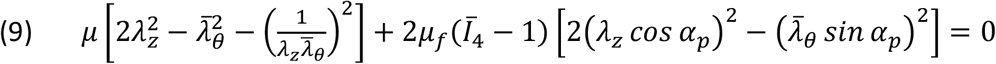

which can be solved for the stiffness ratio between fibers and matrix, *γ*_*f*_ = *μ*_*f*_/*μ*,

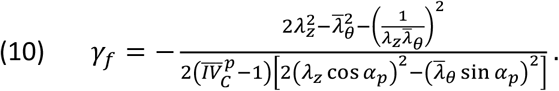

Using (10) one can obtain the required fiber/matrix stiffness ratio for a prescribed value for the orientation of the passive fibers *α*_*p*_ and stretches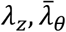.

To quantify the P-V loop in the symmetric design, we follow a similar, albeit simplified, analysis as in the asymmetric case. It is convenient to satisfy equilibrium in three (instead of four in the asymmetric case, again for simplicity) quasi-equilibrated states for the distending pressure (**Figure 3B**): first, *P*_1_ at the end of passive filling (with no myofiber contraction), second, *P*_2_ at the end of isovolumetric contraction with both valves closed, third, *P*_3_ at the end of ejection (with passive deformations coming only from placement pre-stretches) and, fourth, we assume that *P*_4_∼0 at the end of isovolumic relaxation with effectively no contraction (but unlike the asymmetric case, here we do not consider the initial deformation due to the placement pre-load). For the radially averaged assumption, which facilitates intuitive understanding, (7) requires for the three successive states

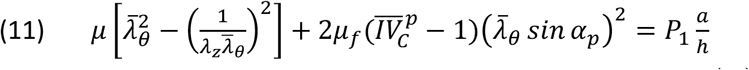

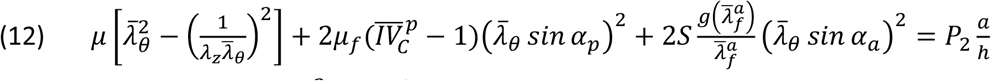

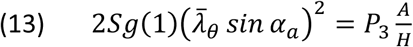

where *g*(1) denotes the value of the activation function 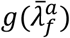 evaluated at 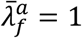 since we assumed that the conduit is undeformed at states 3 and 4.

The solution of (11)-(12) is given by

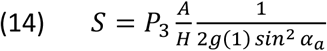

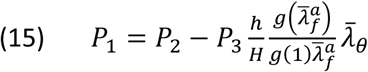

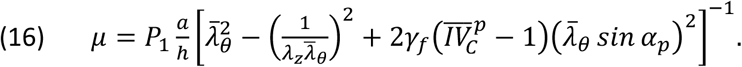

By substitution of the expression for *S* from (14) into (8), the contraction force at the end of isovolumic contraction is given by

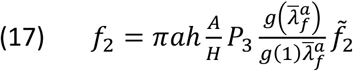

where the normalized force function is given by

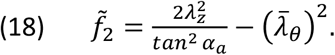

For completeness, we write the expression for EF in the symmetric case including the effect of myofiber deposition stretch λ_0_ (without pre-stretch due to placement)

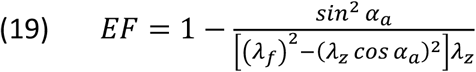

where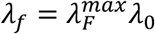.

### Estimation of the relaxed thickness

The circumferential stress from Laplace’s relation for the mean value of the circumferential stress (maximal pressure and homeostatic stress are prescribed) is

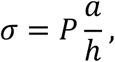

where the luminal radius (with SV, length at end diastolic filling, and EF prescribed) is

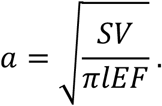

Hence, the loaded thickness in the contracted state is

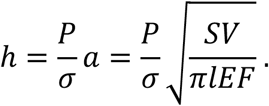

The outer radius when fully loaded (contracted, end-diastolic) is

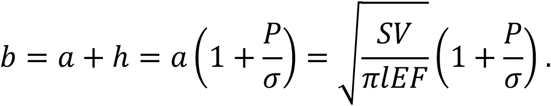

The luminal radius at the relaxed condition (end-systolic), from the relation between EF and biaxial stretches (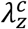 is prescribed), is

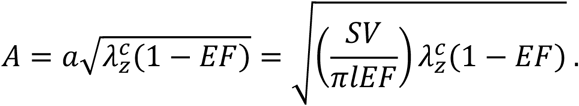

The outer radius at the relaxed condition (from incompressibility) is

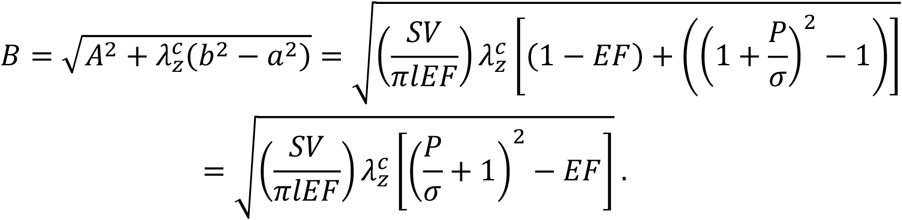

Thus, the relaxed thickness is

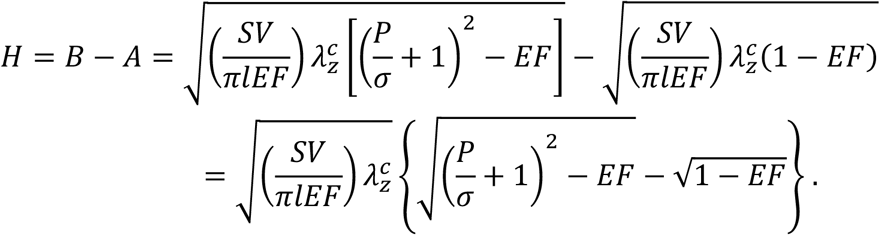

### Species-specific values for the relaxed thickness (H)

Take common values for all three species:

1. 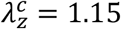
2. *P* = 20 *mmHg* = 2.667 *kPa*
3. *σ* = 30 *kPa* (Burns et al., 1971)
4. *EF* = 0.7

Species-specific values for SV and end diastolic length (Table S2), and the resulting relaxed thickness (H):

|  | Parameter | Human child 3-4 years of age | Sheep | Mouse |
| --- | --- | --- | --- | --- |
| Prescribed | SV (ml) | 17 | 23 | $31 \times 10^{-3}$ |
|  | l (mm) | 50 | 100 | 12 |
| Result | H (mm) | $1.98 \sim 2.0$ | $1.64 \sim 1.7$ | $0.17 \sim 0.2$ |
|  | A (mm) | 7.3 | 6.0 | 0.6 |
|  | B (mm) | 9.3 | 7.6 | 0.8 |
|  | h (mm) | 1.1 | 0.9 | 0.1 |
|  | a (mm) | 12.4 | 10.2 | 1.1 |
|  | b (mm) | 13.5 | 11.1 | 1.2 |

## SUPPLEMENTAL TABLES

**Table S1.** Representative values of key cardiovascular variables for children. Some of the values are based on literature survey: SV (de Simone et al., 1997), EF (Hurwitz et al., 1984), diameter (Itatani et al., 2009), and length (Alexi-Meskishvili et al., 2000), while the remaining can be evaluated via the model equations.

| Parameter | Meaning | Children Range |
| --- | --- | --- |
| $SV$ (ml) | Stroke volume | 15-20 |
| $EF$ (—) | Ejection fraction | 0.5-0.8 |
| $d$ (mm) | End diastolic luminal diameter | 14-25 |
| $l$ (mm) | End diastolic length | 40-60 |
| $D$ (mm) | End systolic luminal diameter | 8-14 |
| $L$ (mm) | End systolic length | 30-50 |
| $\lambda_z$ (—) | Axial cyclic stretch | 1.0-1.2 |
| $\lambda_\theta$ (—) | Circumferential cyclic stretch | 1.0-1.8 |
| $r\varphi$ (—) | Shear deformation | 0.5-1.5 |
| $\Phi$ (deg) | Total twist angle | 0-200 |
| $\alpha_a$ (deg) | Myofiber orientation | -50 to -70 |
| $\alpha_p$ (deg) | Passive fiber orientation | -50 to -70 |
| $\lambda_F^a$ (—) | Myofiber stretch wrt the fiber reference state | 1.0-1.30 |
| $\lambda_f^a$ (—) | Myofiber stretch wrt the tissue reference state | 1.0-1.17 |
| $\lambda_0$ (—) | Myofiber deposition stretch | 0.9-1.00 |
| $\lambda_F^{max}$ (—) | Myofiber maximal stretch | 1.0-1.3 |
| $\lambda_f^p$ (—) | Passive fiber stretch (wrt to the tissue reference state) | >1.0 |

**Table S2.**
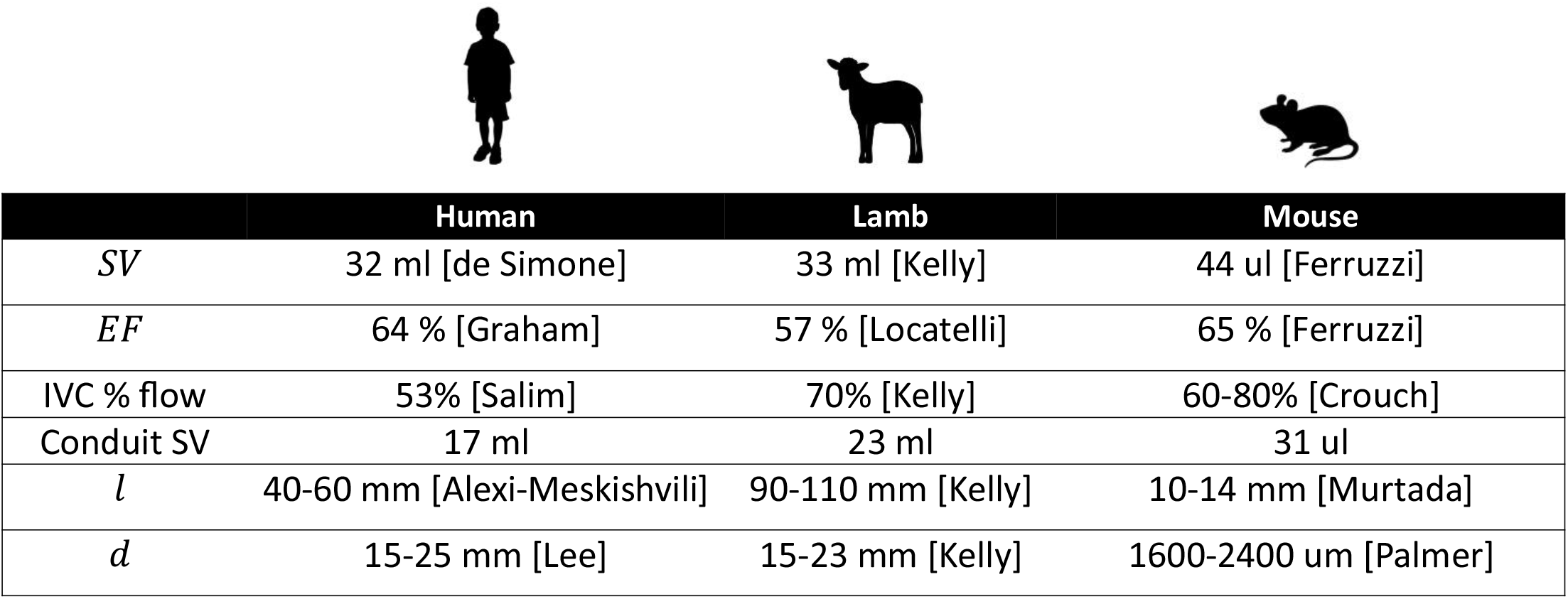
Physiological and geometric parameters for three species (human, sheep, and mouse); see Supplemental References.

|  | Human | Lamb | Mouse |
| --- | --- | --- | --- |
| *SV* | 32 ml [de Simone] | 33 ml [Kelly] | 44 ul [Ferruzzi] |
| *EF* | 64 % [Graham] | 57 % [Locatelli] | 65 % [Ferruzzi] |
| IVC % flow | 53% [Salim] | 70% [Kelly] | 60-80% [Crouch] |
| Conduit SV | 17 ml | 23 ml | 31 ul |
| *l* | 40-60 mm [Alexi-Meskishvili] | 90-110 mm [Kelly] | 10-14 mm [Murtada] |
| *d* | 15-25 mm [Lee] | 15-23 mm [Kelly] | 1600-2400 um [Palmer] |

**Table S3.** Optimized values of key design parameters for three species (human, sheep, and mouse) for the case of unequal orientations (*α*_*a*_ ≠ *α*_*p*_) of active (a) myofibers and passive (p) fibers. The constraint 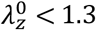 was used in all optimizations. Comparison of values of the objective function between species is meaningless since the range of the force in *J* is different for mouse (*f* ∼ 1 mN) and human/sheep (*f* ∼ 100 mN), also *d*_*max*_ is very different between species.

|  | Parameter | Human | Lamb | Mouse |
| --- | --- | --- | --- | --- |
| Optimized parameters | $\alpha_a$ (deg) | -59 | -60 | -59 |
| | $\alpha_p$ (deg) | -53 | -55 | -54 |
| | $\lambda_F^{max}$ (-) | 1.26 | 1.29 | 1.29 |
| | $\lambda_0$ (-) | 0.92 | 0.91 | 0.91 |
| | $\lambda_z^c$ (-) | 1.13 | 1.13 | 1.13 |
| Preload and pre-stretch | $f_0$ (mN) | 57 | 46 | 0.49 |
| | $\lambda_z^0$ (-) | 1.13 | 1.16 | 1.15 |
| | $\bar{\lambda}_\theta^0$ (-) | 1.00 | 0.99 | 0.99 |
| Myofiber and passive stretch at end-diastole | $\lambda_F$ (-) | 1.24 | 1.25 | 1.25 |
| | $\lambda_f^p$ (-) | 1.01 | 1.01 | 1.01 |
| Material parameters | $\mu$ (kPa) | 1.51 | 1.43 | 1.33 |
| | $\gamma_f$ (-) | 8.9 | 9.1 | 8.5 |
| | $S$ (kPa) | 31.11 | 30.68 | 27.46 |
| Unloaded geometry | $L_0$ (mm) | 39.3 | 75.9 | 9.3 |
| | $R_0$ (mm) | 8.2 | 6.9 | 0.74 |
| | $H_0$ (mm) | 2.2 | 1.95 | 0.23 |
| Peak loads | $f_{max}$ (mN) | 79 | 51 | 0.70 |
| | $T_{max}$ (mN × mm) | 1.82x10<sup>4</sup> | 1.01x10<sup>4</sup> | 12.14 |
| Objective function | $J(-)$ | 5.4554 | 2.4659 | 3.2869e-04 |

**Table S4.** Ranges of values of key design parameters for three combinations of myofiber shortening and deposition stretch shown in Figure 7. Case B had the least optimized loads, namely, the highest range of force and torque, but the largest admissible design space. In contrast, Case C had the best optimized loads, namely, the lowest range of force and torque, but with a tighter design space. Also note that the reserve in myofiber stretch is minimized in Case C, namely the lower end of the range for λ_*F*_ is closer to 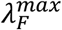, whereas in Cases A and B the reserve in myofiber stretch is greater, reflecting poor use of myofiber contractility.

| Case | $\lambda_F^{max}$ (—) | $\lambda_0$ (—) | $\vert T_{max}\vert $ ($mN \times mm$) | $f_{max}$ ($mN$) | $\lambda_F$ (—) |
| --- | --- | --- | --- | --- | --- |
| A | 1.2 | 0.95 | $[1.77, 2.35] \times 10^4$ | [216, 912] | [1.08, 1.20] |
| B | 1.3 | 0.95 | $[1.78, 2.70] \times 10^4$ | [221, 1573] | [1.08, 1.30] |
| C | 1.3 | 0.90 | $[1.63, 1.76] \times 10^4$ | [78, 296] | [1.21, 1.30] |

## SUPPLEMENTAL FIGURES

**Figure S1:**
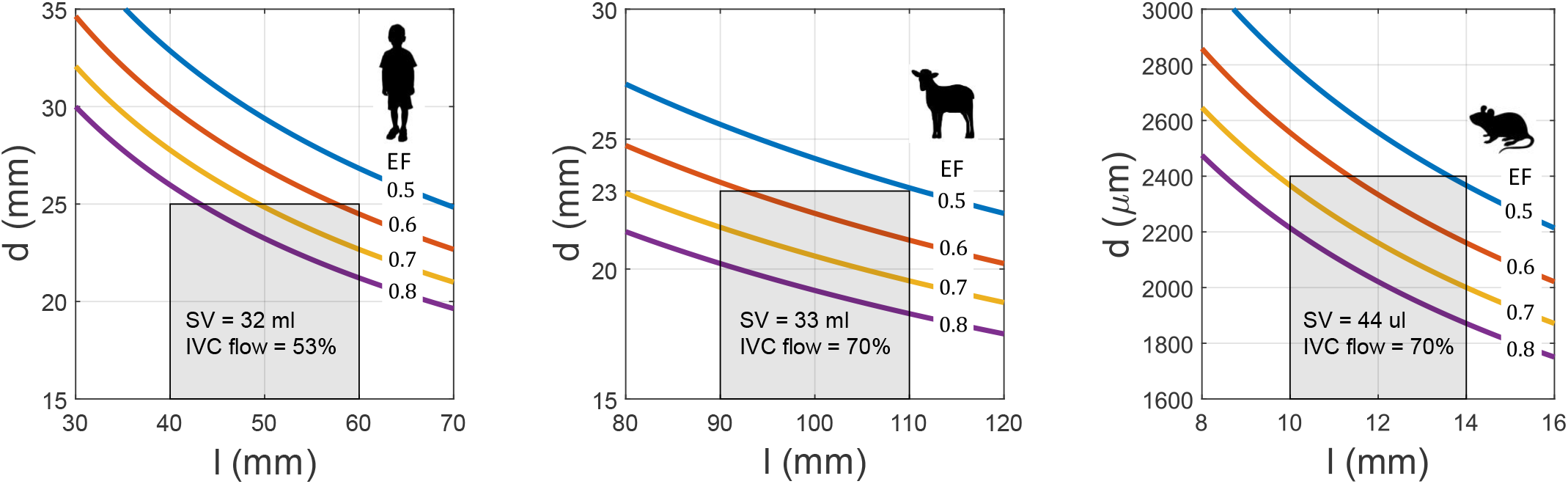
Similar to Figure 1B but across three species (the human panel is redundant but included for direct comparison). The parameter values used to generate these figures are in Table S2 (maximal luminal diameter and length, SV, EF).

**Figure S2:**
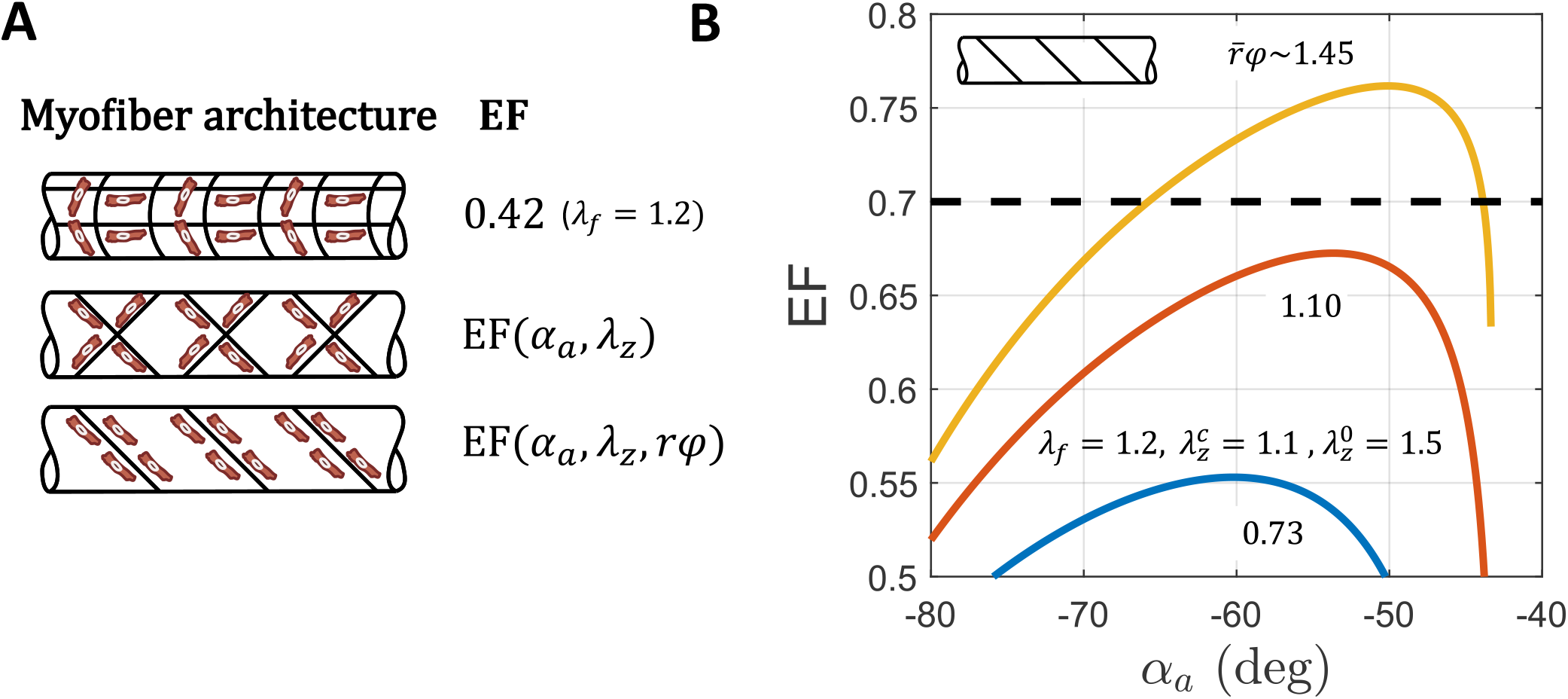
Similar to Figure 2, except with less robust myofiber shortening (λ_*f*_ = 1.2).

**Figure S3:**
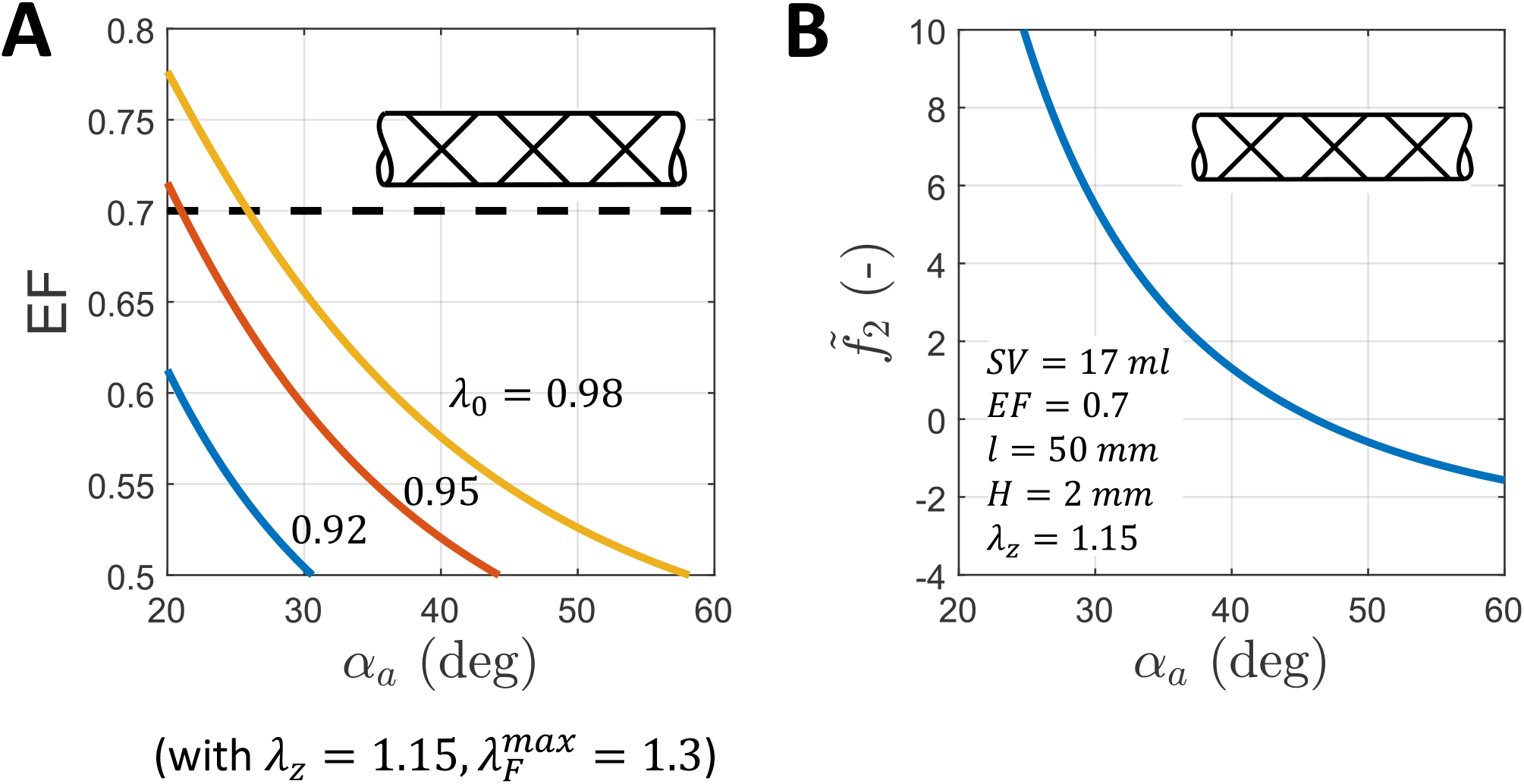
Parametric results for a symmetric orientation of the myofibers. (A) Ejection fraction and (B) normalized axial load. Supplementary equations (19) and (18) were used to generate the specific examples for EF and normalized axial force 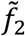 at end isovolumic contraction, respectively. In panel A, values for *EF* > 0.7 are available only for a limited range of active myofiber orientations *α*_*a*_∼30 deg, even with almost no deposition stretch (λ_0_ → 1). In panel B, normalized force close to zero corresponds with orientation *α*_*a*_∼45 deg that falls below the *EF* = 0.7 threshold. However, active myofiber orientations with *EF* = 0.7 are aligned more to the axial direction and generate higher axial force than in the asymmetric diagonal fiber architecture (compare with Figure 4A). Although not shown here, the filling pressure *P*_1_ (easily calculated from supplementary equation (15)) has negative values for realistic values of deposition stretch λ_0_∼ 0.92 – 0.95, and myofiber orientations *α*_*a*_∼ 20 – 40 deg, which is unwanted.

**Figure S4:**
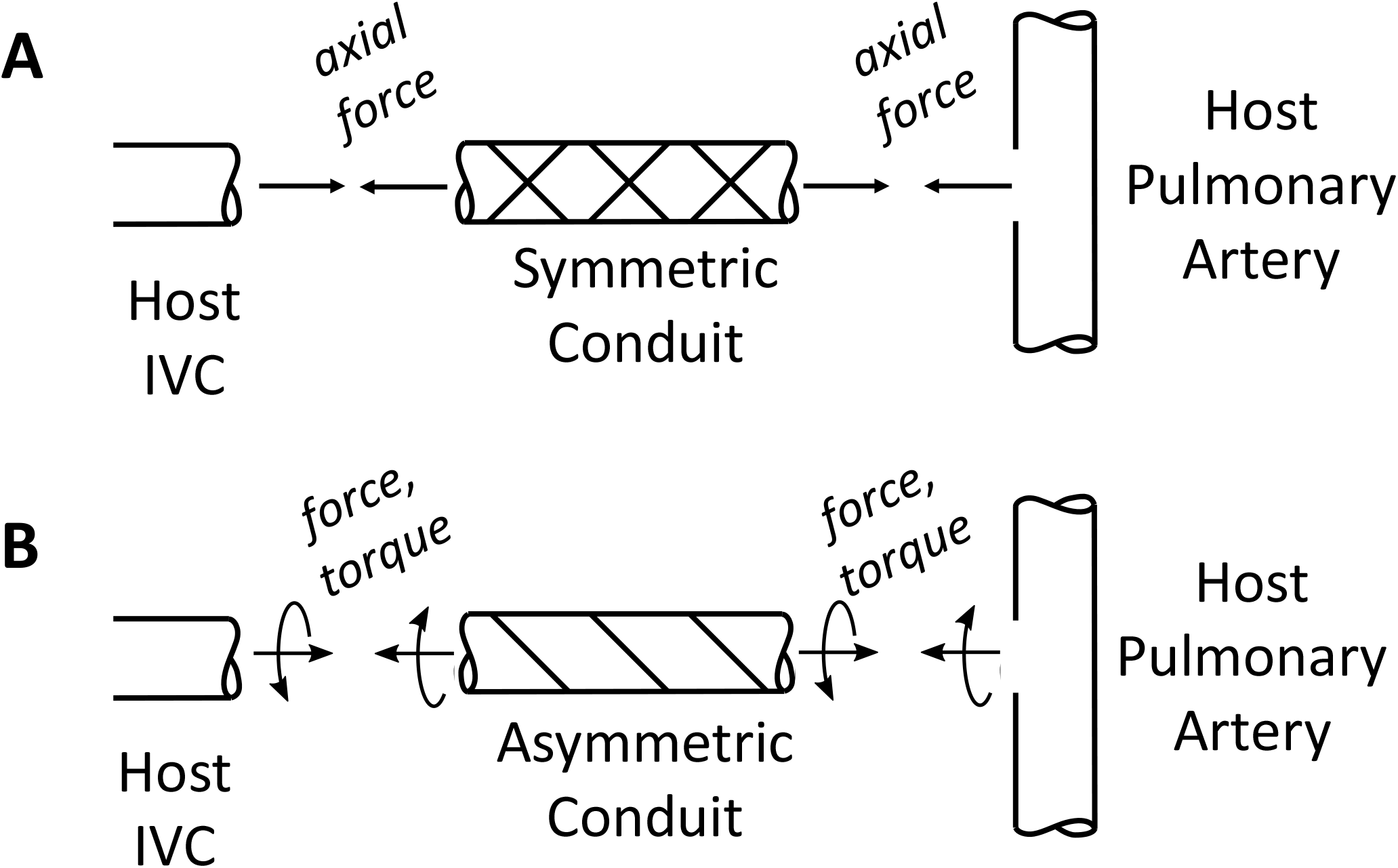
Schema emphasizing that, by Newton’s 3^rd^ law of motion, (A) pulsatile conduits with dual families of symmetric diagonal myofibers will generate axial forces at the anastomoses while (B) conduits with single families of asymmetrically oriented myofibers will generate axial forces and torques at the anastomoses.

**Figure S5:**
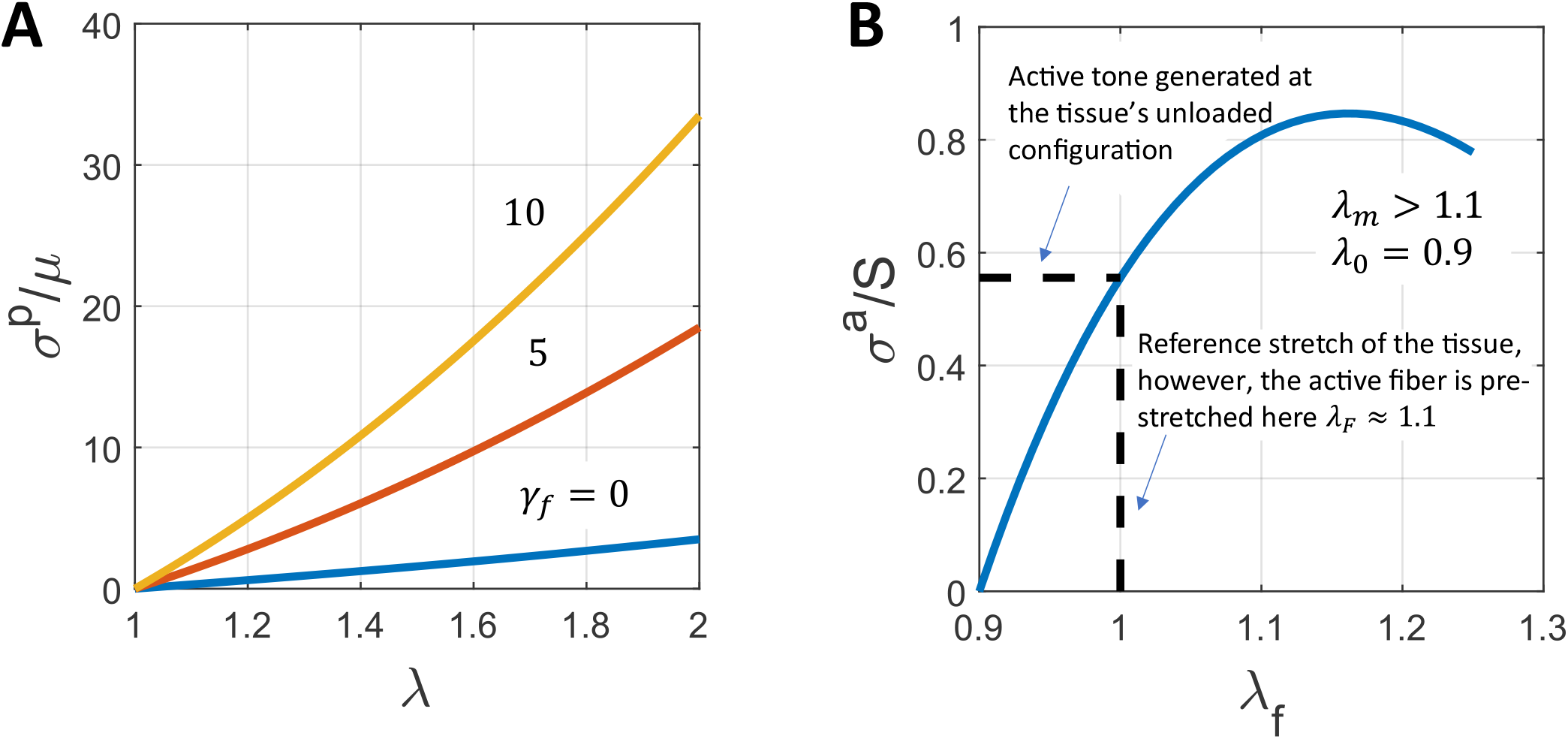
Normalized uniaxial Cauchy stress for (A) passive and (B) active constituents (based on constitutive relations in Equations (12) and (13), respectively). Panel A shows effects of different levels of passive fiber reinforcement (*γ*_*f*_ = 0, 5, 10) where *γ*_*f*_ = *μ*_*f*_/*μ* denotes the passive fiber stiffness relative to an amorphous matrix stiffness (with zero denoting no fibers). Panel B shows a representative active stress behavior (normalized). The stress generated depends on the level of myofiber shortening from stretch λ_*f*_, which is defined with respect to overall tissue deformation with λ_0_ a “ deposition stretch” at which force generation is zero. This curve recapitulates a Frank-Starling behavior with λ_*f*_ = λ_*m*_ denoting the maximal normalized Piola–Kirchhoff stress.

**Figure S6:**
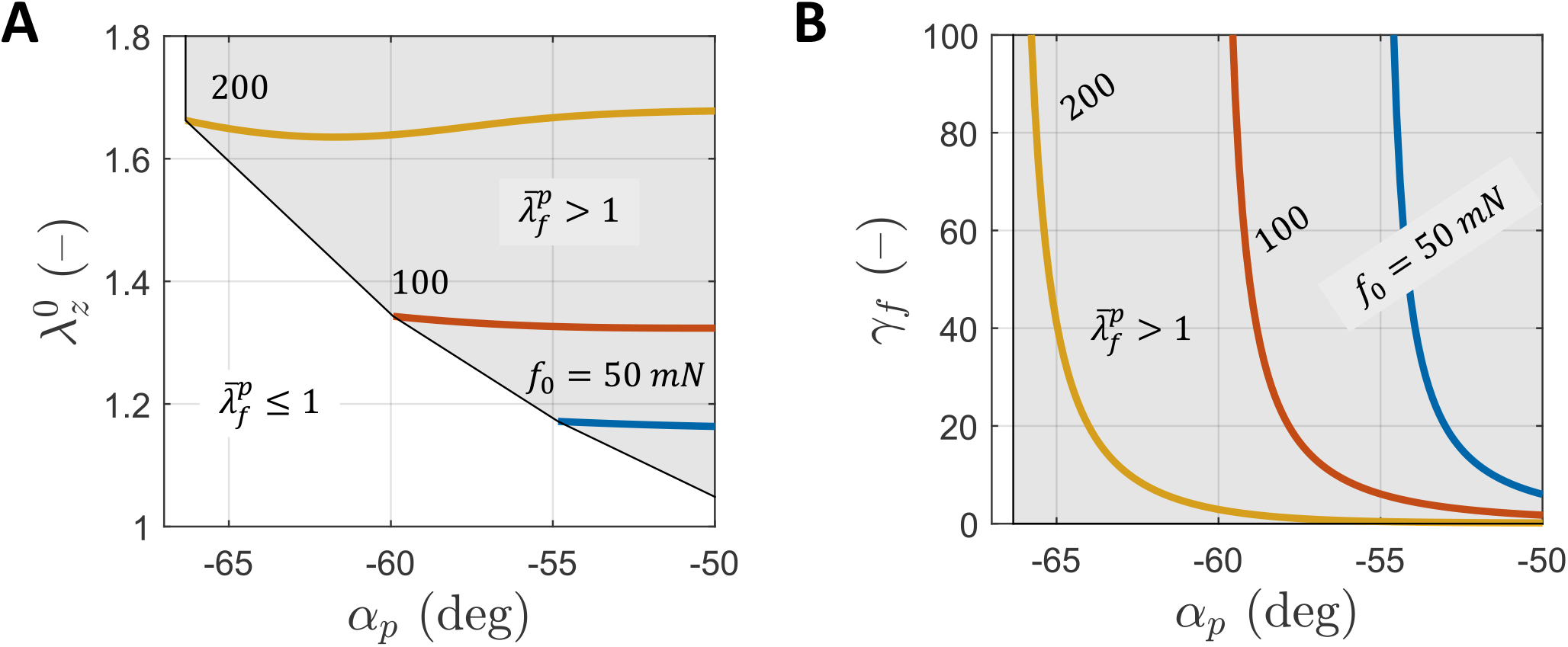
Additional effects (compare with Fig. 4A) of passive fiber orientation (via angle *α*_*p*_) on (A) the axial pre-stretch at implantation 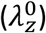 and (B) the relative fiber stiffness with respect to matrix stiffness (*γ*_*f*_) as a function of three different values of the initial axial force (*f*_0_ = 50, 100, 200 mN). The grey zones show admissible values for which the passive fibers remain in tension 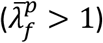 whereas white zones exclude inadmissible values 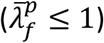. Note that 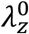 is nearly independent of passive fiber orientation (beneficial from biomanufacturing perspective), and that *γ*_*f*_ asymptotes near a threshold orientation, depending on the value of the initial axial force. This specific example was generated by solving equations (30)-(33) for 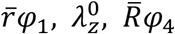, and *γ*_*f*_ for a range of passive fiber orientations *α*_*p*_ while prescribing representative values for *μ* = 1 kP, 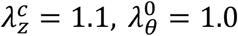. Other parameters values used to generate these figures are SV = 17 ml, EF = 0.7, d = 25 mm, l = 50 mm, H = 2 mm.

**Figure S7:**
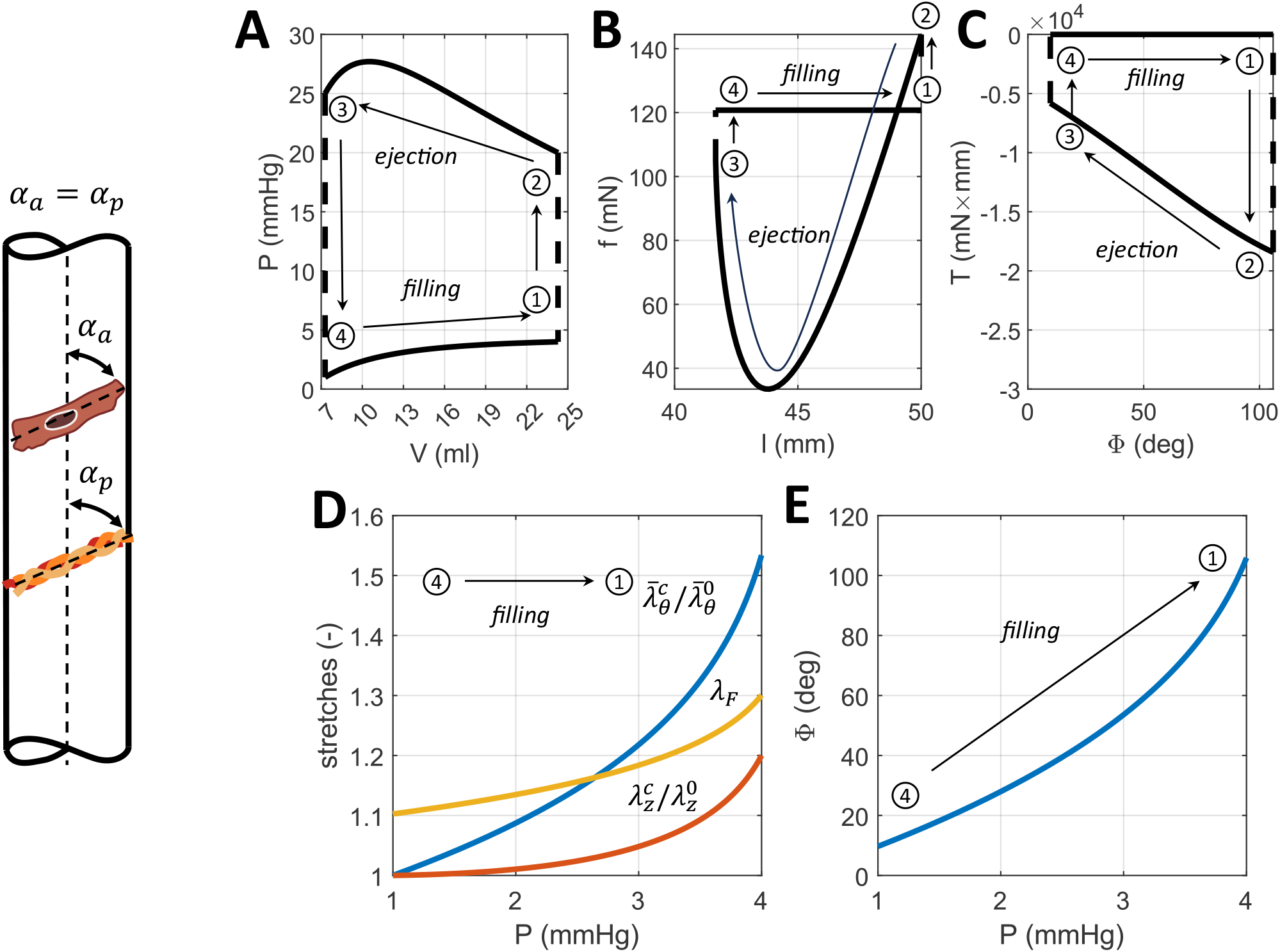
The same as Figure 6 except for optimized values of the design parameters for equal active myofiber and passive fiber orientations (*α*_*a*_ = *α*_*p*_) under the constraint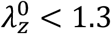. See also Table 1.

**Figure S8:**
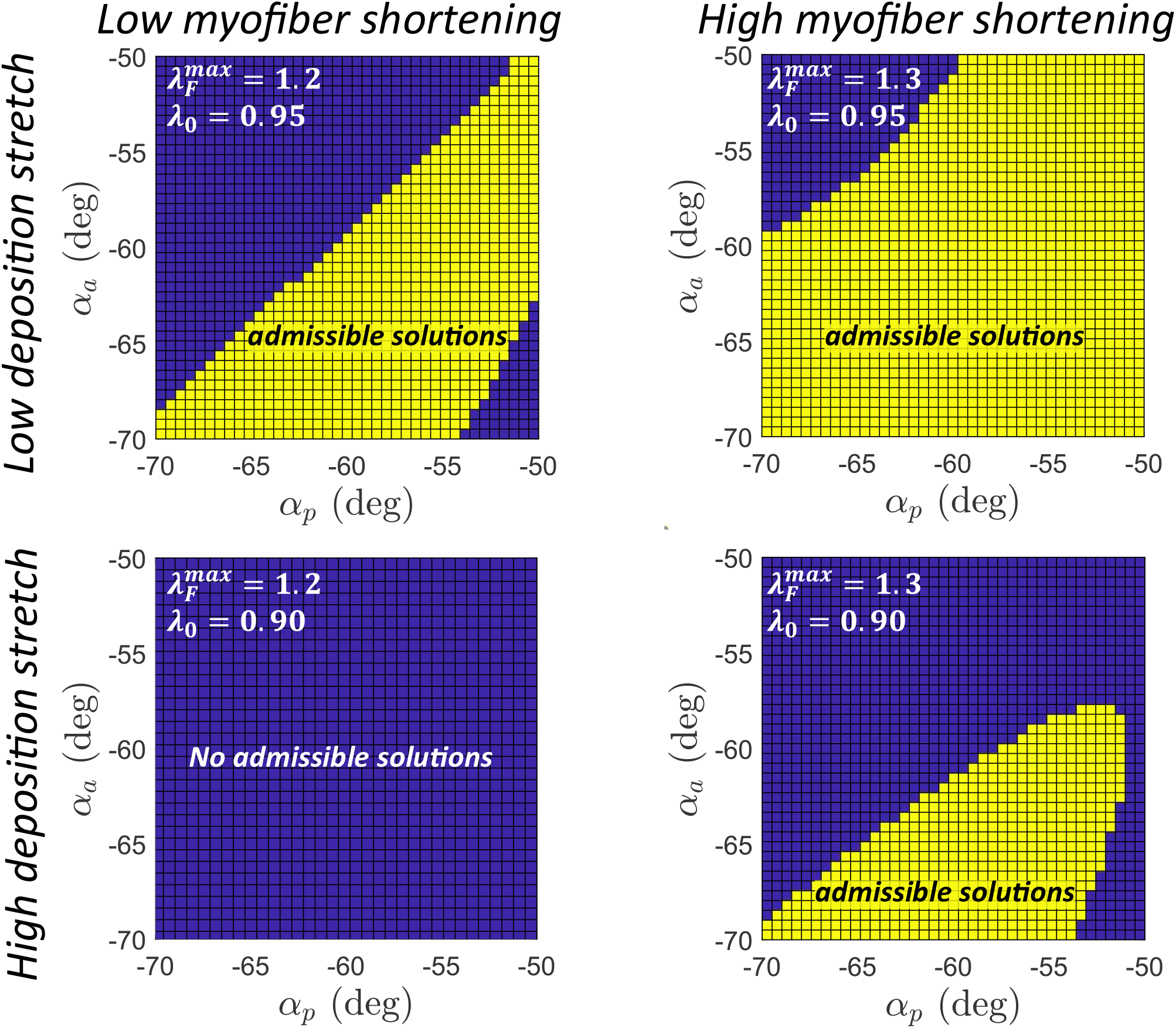
Mathematically admissible parameter spaces for different combinations myofiber shortening (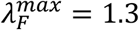 and 1.2, for high and low, respectively) and myofiber deposition stretch (λ_0_ = 0.90 and 0.95, for high and low, respectively). The parameters values used to generate these figures are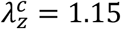, SV = 17 ml, l = 50 mm, H = 2 mm, EF = 0.7, P4 = 1 mmHg, P1 = 4 mmHg, P2 = 20 mmHg, P3 = 25 mmHg. Note that not all mathematically admissible solutions yield a biomechanically valid solution (see Figure 7).

**Figure S9:**
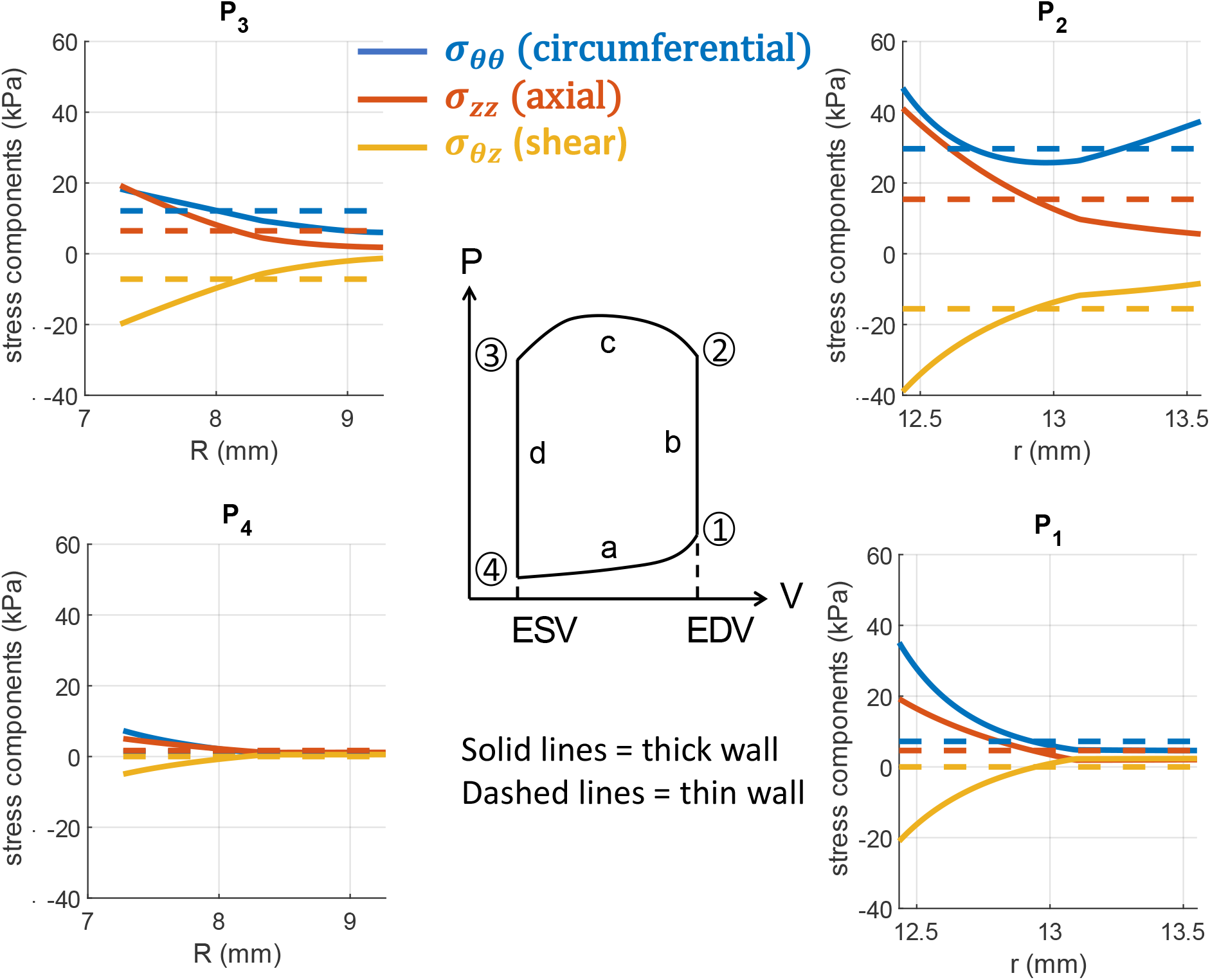
Illustrative comparison of potential transmural distributions of multiaxial wall stress (circumferential – blue, axial – red, and shear – yellow) at the different pressures along a pressure-volume loop (center, P1-end of passive filling, P2-end of isovolumic contraction, P3-end of ejection, P4-end systole). Note that the highest wall stresses are at P2. Dashed lines show radially averaged approximations, solid lines show computed thick-walled values in the absence of residual stress.

**Figure S10:**
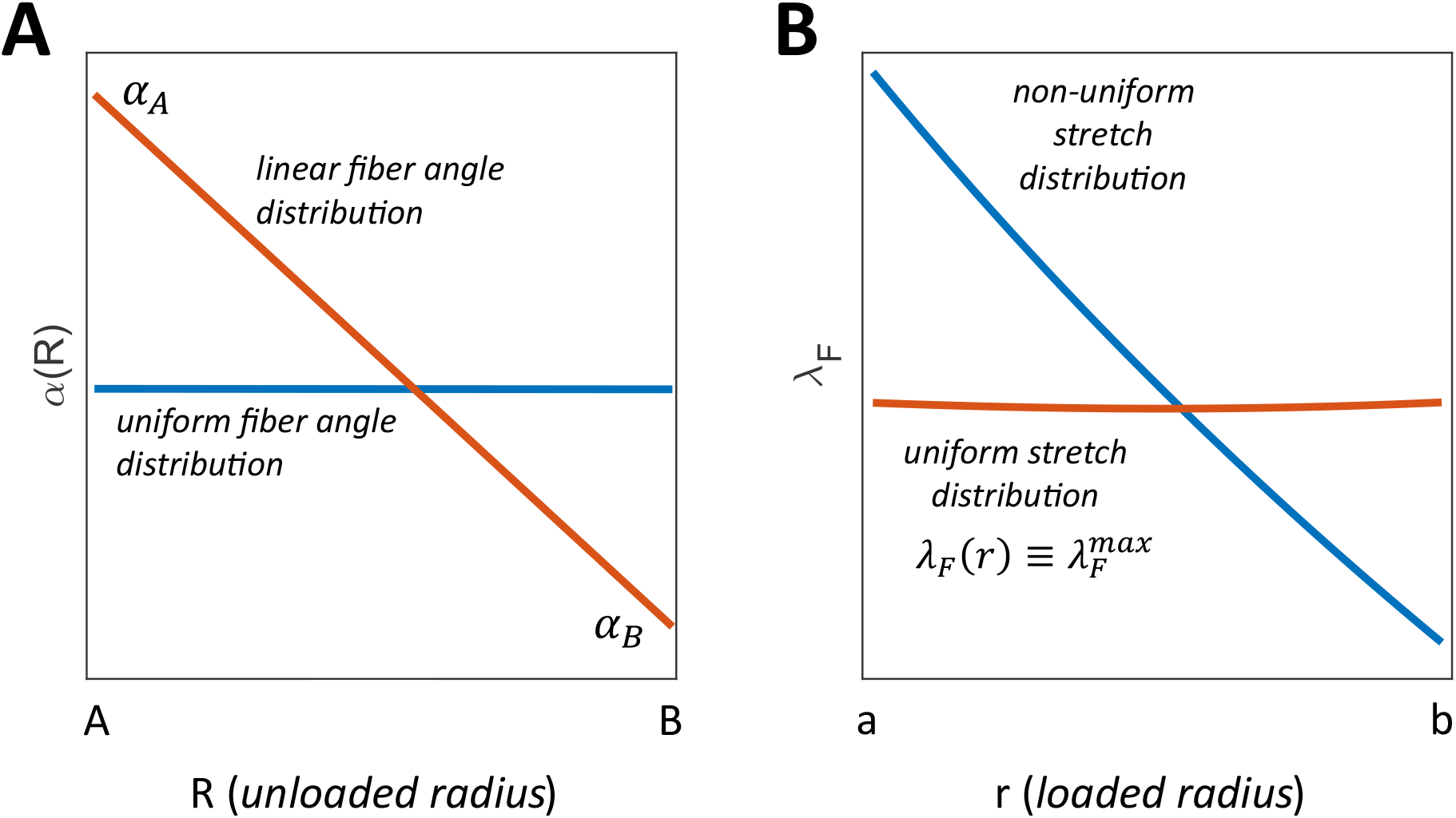
Qualitative demonstration that a linear transmural splay of myofiber directions through a thick-walled conduit (orange - left) can yield a uniform distribution of myofiber stretches (orange - right), which appears to be most efficient. By contrast, a uniform value of fiber orientations through the wall (blue – left) would yield a transmural distribution of myofiber stretches (blue – right) that would be expected to be mechanobiologically unfavorable. Note that the former is inspired by transmural distributions of myofiber orientation in the native ventricle. It is anticipated that conduits having thicker walls may need to have nonlinear distribution of myofiber orientations. It is also possible that radial variations in other properties like matrix shear modulus, fiber stiffness, active stress, and orientation of the passive matrix (*μ, μ*_*f*_, *S, α*_*p*_, respectively), along with residual stresses, can further optimize the stress distribution.

## Notes

### Competing Interest Statement

The authors have declared no competing interest.

